# Horizontal gene transfer fuels metabolic innovation in the grass *Zuloagaea bulbosa*

**DOI:** 10.64898/2026.06.09.731121

**Authors:** Lara Pereira, Emily J Bailes, Noah G Bourne, Catherine F Collins, Sahr Mian, Ilia J Leitch, Benjamin R Lichman, Luke T Dunning

## Abstract

Horizontal gene transfer (HGT) allows the movement of DNA across broad evolutionary distances without sexual reproduction. In grasses, HGT is widespread and although a few horizontally transferred genes (HTG) are adaptive, most are purged over time. Within the adaptive HTG, biosynthetic genes encoding enzymes that act together in the same pathway and physically co-localise in clusters have been reported multiple times. The aims of this study are to test whether HGT is bidirectional in a pair of grass species, maize and *Zuloagaea bulbosa*, and if HTGs are found more than expected by chance in biosynthetic genes organised in clusters. To achieve this, we firstly generated a phased reference genome for *Z. bulbosa*. Then we identified 56 candidate horizontally transferred genes, of which 45% were from Andropogoneae, including two likely to be of maize origin. Since transfers from *Z. bulbosa* to maize were previously described, our results show that HGT is bidirectional, although the balance might not be even. After predicting all biosynthetic gene clusters in the *Z. bulbosa* genome, we found that HTGs are enriched in biosynthetic genes organised in clusters. This correlation between HGT and gene clustering is likely to be a consequence of selection due to the immediate adaptive benefit a whole pathway can provide. Two of the HTGs from Andropogoneae belong to the benzoxazinoid BGC, which previously underwent an ancestral transfer from Panicoideae into Pooideae. The dynamism of biosynthetic gene clusters, including recurrent horizontal gene transfers, contributes to the extraordinary metabolic diversity present in plants.

**Significance statement:** Horizontal gene transfer (HGT) is a significant driver of evolution that is widespread in grasses. In this study, we show for the first time reciprocal transfer of DNA between maize and another Mexican grass, *Zuloagaea bulbosa*. This result represents a proof of concept of bidirectional HGT which allows for limited, recurrent gene flow among distant species. Furthermore, we show that horizontally transferred genes are enriched for biosynthetic genes organised in biosynthetic gene clusters, regions of the genome that encode for multiple enzymes that act in the same biosynthetic pathway. We hypothesise that the transfer of a complete multi-genic pathway, ready to be used and potentially offering an evolutionary advantage, might promote a predominant retention of gene clusters in comparison with background HGT.

## Introduction

Gene flow across species boundaries can have a longstanding impact in evolution and adaptation (Ellstrand, 2014; Ellstrand & Rieseberg, 2016; X. Huang et al., 2025; Tigano & Friesen, 2016). Hybridisation, defined as interbreeding of two individuals from distinct species, is common in plants and has immediate phenotypic consequences (Harrison & Larson, 2014; Whitney et al., 2010). While hybridisation is common between closely related taxa, horizontal gene transfer (HGT) can also move trace amounts of DNA across broader evolutionary distances by means other than sexual reproduction.

Plant-to-plant HGT has been increasingly reported over the last few decades as more reference genomes became available (Christin et al., 2012; F.-W. Li et al., 2014; D. Wu, Hu, et al., 2022; D. Wu, Jiang, et al., 2022; Yang et al., 2016, 2019; Zhang et al., 2014). Parasitic plants live in physical proximity with their hosts and are able to acquire large genomic fragments from them (Yoshida et al., 2010, 2019). This transfer is unidirectional from host to parasite, and close physical interactions via a specialised structure called the haustorium are thought to facilitate the movement of genetic material (Yoshida et al., 2019). Outside of parasitic plants, HGT seems to be prevalent in grasses (Hibdige et al., 2021; Y. Huang et al., 2026; Mahelka et al., 2017, 2021; Park et al., 2021), where many HGT events are neutral or deleterious and purged over time (Raimondeau et al., 2023), but a proportion of HGTs are maintained and selected for (Christin et al., 2012; Olofsson et al., 2019). Large genomic fragments (e.g. >200 kb) can be transferred (Dunning et al., 2019; Mahelka et al., 2021), including those containing multiple protein-coding genes with their cis-regulatory elements (Collins et al., 2026). Therefore, it is now established that plant-to-plant HGT can be an important driver of adaptation and genome evolution (Van Etten & Johnson, 2026).

Donor-recipient dynamics, especially in recent, intra-family HGT events, are key to understanding the mechanisms behind DNA movement and the ecology of HGT beyond genome integration (Van Etten & Johnson, 2026). Phylogenetic methods estimate likely donor clades, with support from biogeographical patterns. However, identifying the exact donor of a given horizontally transferred gene (HTG) is difficult, since most transfers happened several million years ago and were acquired from the common ancestors of extant species (Raimondeau et al., 2023). Despite these challenges, studying donor-recipient dynamics with comparative genomic methods is paramount. By comparing the genomes of two species that have recently exchanged DNA we can see how fragments erode over time, and how gene regulation and even gene function evolve when integrated in a novel genomic context. Of special interest is to identify patterns of bidirectional transfer between pairs of species, since this genetic dialog could facilitate co-evolution. Outside of parasite-host interactions, reciprocal HGT has not been studied widely in plants.

Maize (Zea mays, Andropogoneae) has huge agricultural relevance and a well-studied genome (Hufford et al., 2021; Schnable et al., 2009). The maize genome carries 11 horizontally acquired genes, half of them acquired from the subtribe Cenchrinae (Paniceae) (Hibdige et al., 2021). Of the HTGs in maize that were acquired from Cenchrinae, the species *Zuloagaea bulbosa* appeared as a sister to three of them (∼30% of total HTGs) (Fig. 1) (Hibdige et al., 2021). *Z. bulbosa* is a wild, weed-like species found in the south of North America, mainly in Mexico and the south of the United States (Bess et al., 2006), which is where maize was domesticated 9,000 years ago (Stitzer & Ross-Ibarra, 2018). Given the phylogenetic and biogeographic context, we hypothesised that the *Z. bulbosa* lineage recurrently acted as donor for horizontally acquired genes in maize, making this pair of species a useful system to study HGT dynamics post-transfer.

**Figure 1.**
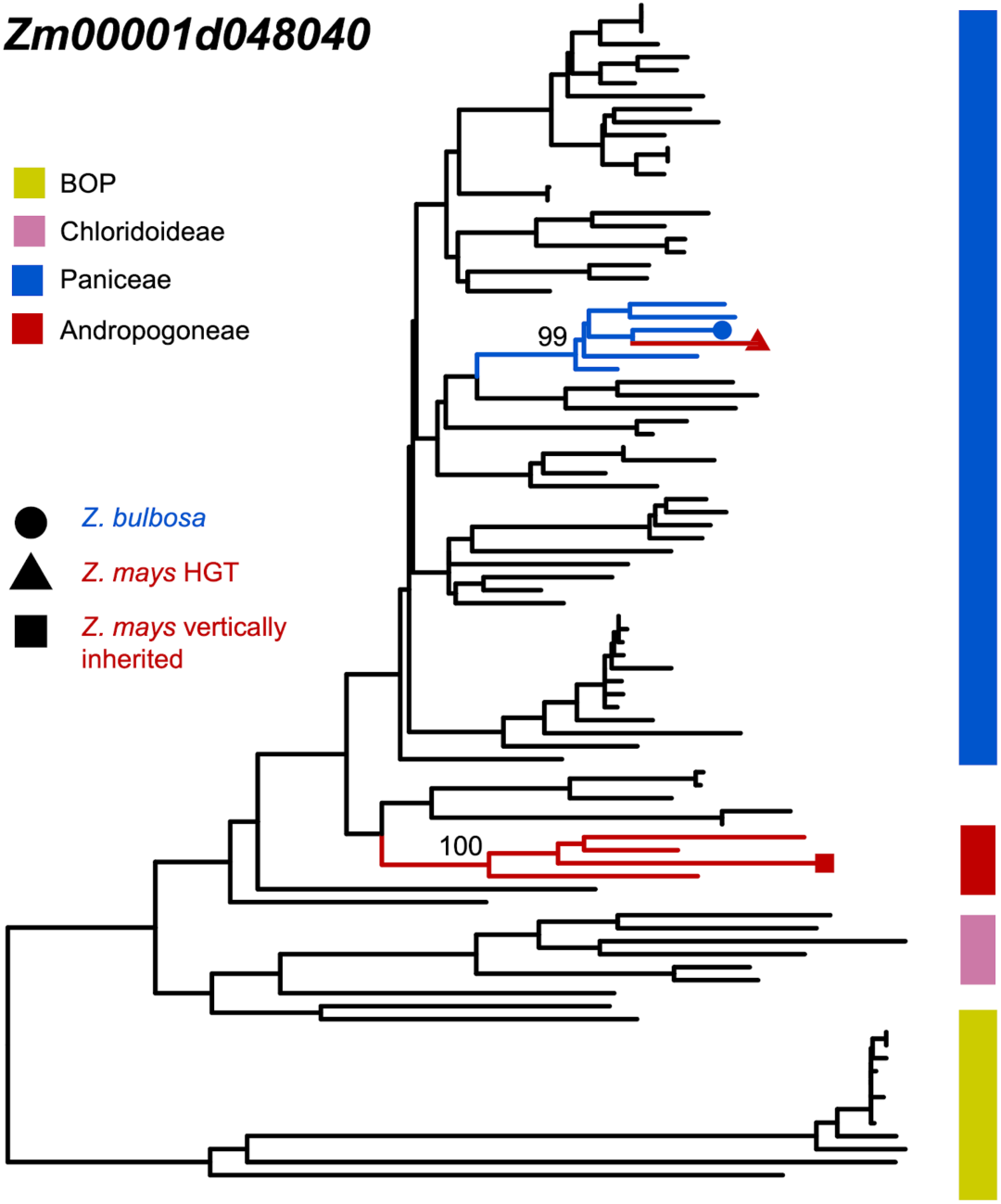
One representative example of the three maize HTGs nested in Cenchrinae, and sister to *Z. bulbosa* ortholog. In the tree, Cenchrinae clade is represented in blue and Andropogoneae, in red. The tree has been redrawn from Hibdige et al. (2021).

Among the few well-characterised examples of HTGs in grasses, several involve biosynthetic gene clusters (BGCs) (Y. Huang et al., 2026; D. Wu, Hu, et al., 2022; D. Wu, Jiang, et al., 2022). BGCs are co-localising, tightly linked non-homologous genes that encode different enzymes from the same biosynthetic pathway (Nützmann et al., 2016; Smit & Lichman, 2022). BGCs are largely located in highly dynamic genomic regions (Field et al., 2011), seem to undergo frequent genomic rearrangements (Liu et al., 2023), possibly mediated by miniature inverted-repeat transposable elements (Boutanaev & Osbourn, 2018). The process through which biosynthetic genes assemble into clusters is currently unknown, despite several hypotheses recently proposed (Cawood & Ton, 2024; Smit & Lichman, 2022). Previous research has shown that the momilactones, benzoxazinoids, and gramine gene clusters have been horizontally transferred between distant grasses (Y. Huang et al., 2026; D. Wu, Hu, et al., 2022; D. Wu, Jiang, et al., 2022), enabling inter-species exchange of complete metabolic pathways which encode for the biosynthesis of a compound with adaptive benefits. We hypothesise that HGT might also contribute to the assembly and evolution of the BGCs themselves, leading to cluster expansion, contraction and variation.

The main aim of this study was to investigate whether HGT is reciprocal, with the same pair of species acting as either donor or recipient of horizontally transferred DNA fragments. Given the transfer of several BGCs in other grass species, we also tested whether HGT was predominantly observed in genes located within BGCs. To achieve this, we first generated a reference genome for *Z. bulbosa* and identified candidate HTGs. We identified two HTGs which are likely of maize origin, implying that HGT is not unidirectional in this pair of species. We also show that biosynthetic genes organised in clusters are transferred more frequently than expected, and that HGT contributes to BGC evolution and dispersal.

## Results

### A well-resolved *Zuloagaea bulbosa* genome

The 1C holoploid genome size of *Z. bulbosa* (i.e. the amount of DNA in an unreplicated nucleus, regardless of ploidy level) was estimated to be 1.81 Gb by flow cytometry. We obtained 67.57 Gb of HiFi data (5.54 million reads with a mean length of 12.2 kb) and 150 Gb of Hi-C data. Although the k-mer spectrum was best fit by a tetraploid model, the flow cytometric 1C genome size of ∼1.8 Gb is approximately three times the GenomeScope haploid estimate (∼0.62 Gb) (Fig. S1). This suggests that the sequenced individual may be a higher polyploid, potentially hexaploid or a complex segmental polyploid, rather than a simple tetraploid, in agreement with previous studies (Doust et al., 2007).

The hifiasm pipeline yielded 8,339 primary unitigs, comprising a total genome length of 3.167 Gb, which is close to the estimated 2C value (3.62 Gb). Despite using unitigs instead of contigs to avoid collapsing different haplotypes, the total genome size was slightly lower than the genome size estimated by flow cytometry, indicating that some unitigs are likely representing multiple identical haplotypes. We then used Hi-C reads to scaffold the unitigs, improving the contiguity of the assembly (Table 1). The scaffolded assembly had an L90 of 89. After excluding the organellar contigs (1,066 small scaffolds, ranging from 16,023 to 95,566 bp), L90 was reduced to 82 scaffolds. Since this number is only slightly higher than the expected chromosome number if hexaploid (6n≈54-70) (Doust et al., 2007), some of the scaffolds are likely full chromosomes. This is also suggested by whole-genome alignments of *Zuloagaea bulbosa* scaffolds against chromosomes of the Cenchrinae grass *Setaria viridis*, which shows broad conservation of genomic collinearity between the two species (Fig. S2).

**Table 1.**
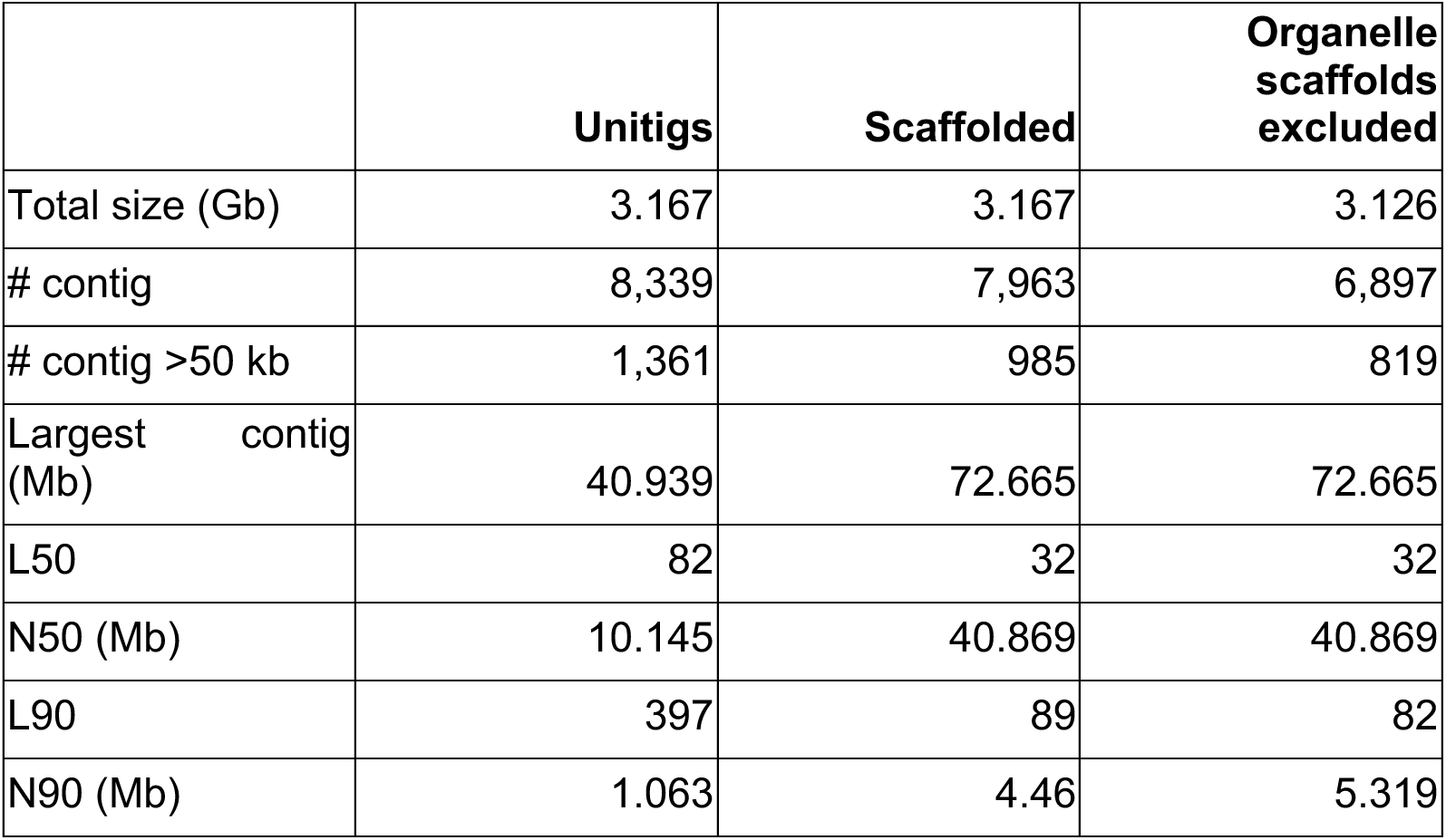
Statistics of *Zuloagaea bulbosa* phased assembly, before and after scaffolding with Hi-C data, and after excluding organelle scaffolds.

|  | Unitigs | Scaffolded | Organelle scaffolds excluded |
| --- | --- | --- | --- |
| Total size (Gb) | 3.167 | 3.167 | 3.126 |
| # contig | 8,339 | 7,963 | 6,897 |
| # contig >50 kb | 1,361 | 985 | 819 |
| Largest contig (Mb) | 40.939 | 72.665 | 72.665 |
| L50 | 82 | 32 | 32 |
| N50 (Mb) | 10.145 | 40.869 | 40.869 |
| L90 | 397 | 89 | 82 |
| N90 (Mb) | 1.063 | 4.46 | 5.319 |

Genome annotation yielded 235,246 gene models, which is ∼6 times of what we would expect in a haploid genome assembly (∼35,000 – 40,000 genes). Completeness was very high (99.8% for the genome assembly and 99.9% for the annotation of Poales BUSCO genes), as well as duplication (99% and 99.1% duplicated respectively), indicating a high-quality assembly.

### Reciprocal horizontal gene transfer between maize and *Zuloagaea bulbosa*

*Z. bulbosa* was previously identified as a likely donor for ∼30% (3 of the 11 maize HTGs) of the horizontally transferred genes found in maize, since it appears as sister to the candidate horizontally transferred genes (HTGs) (Fig. 1) (Hibdige et al., 2021). With the aim of testing if this DNA transfer could be reciprocal, we identified genes of foreign origin in the genome of *Z. bulbosa*. Our phylogenetic pipeline for HGT identification yielded 56 HTG candidates (Table S2). After manual inspection, 31 of these candidates were classified as robust HTGs. For all the robust candidate HTGs, the donor species belong to either the Andropogoneae tribe (45.2%) or the Chloridoideae subfamily (54.8%) (Dataset S1). More than half of the HGT events likely happened before the *Z. bulbosa* polyploidization event(s), since multiple copies of horizontally acquired genes in different scaffolds were found. However, we also found single-copy candidate HTGs, which could be a product of recent events, or alternatively, be due to gene losses post-transfer.

Among the robust HTGs, we found two HGT events for which maize was the likely donor. One of the candidates had many copies from *Z. bulbosa* present in the tree (>100) and these were homologous to a Helitron DNA helicase (HGT_044, Fig. 2A). This Helitron DNA helicase is absent from most grass species (Dataset S1) and the gene tree does not follow the same topology as the species tree. However, all *Z. bulbosa* copies were nested within maize sequences with 100 bootstrap support, and the branch lengths were considerably shorter than those in the rest of the tree, likely indicating recent HGT of a transposable element followed by a burst of transposition in the recipient genome.

**Figure 2.**
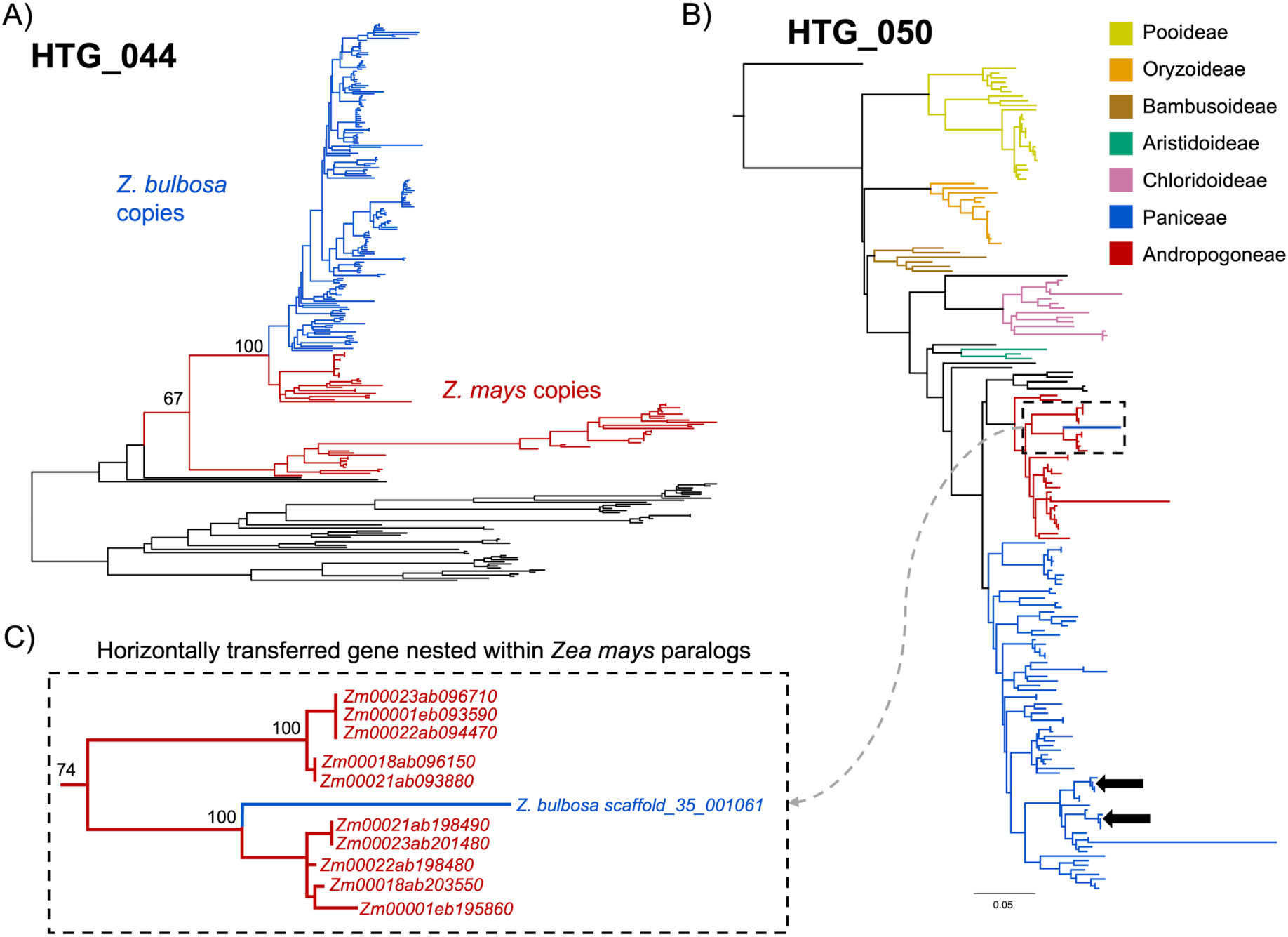
HGT events showing *Z. mays* as likely donor to *Z. bulbosa* HTG. A) Phylogenetic tree for the Helitron DNA helicase (candidate HGT_044). *Z. bulbosa* copies are drawn in blue and *Z. mays,* in red. Bootstrap support indicated for the two nodes showing that *Z. bulbosa* sequences are nested within *Z. mays*. B) Phylogenetic tree for candidate HTG_050, where the black arrows point to *Z. bulbosa* vertically inherited copies and the dashed rectangle encloses the HGT. C) Zoom view into the *Z. mays* clade, which includes two maize paralogs and the HTG from *Z. bulbosa* nested within them.

The other transfer moved three candidate HTGs (HTG_012, HTG_050, and HTG_037) as part of the same DNA fragment. The three genes are single-copy, and appear nested within two maize paralogs (Fig. 2B and C shows one example), indicating that the donor is maize or a closely related species. The presence of this fragment in *Z. bulbosa* genome, together with the Helitron DNA helicase, indicates that HGT can be reciprocal, allowing a limited gene flow between distant grass species that co-exist in the same habitat.

### Horizontally transferred genes are located in biosynthetic gene clusters more than expected by chance

We performed a blast search of all of the HTG candidates against the SwissProt database to obtain a tentative functional annotation. We found that HTG candidates were predicted to be involved in diverse biological functions, such as defence against pathogens and stresses (20.75%), protein quality control (9.43%), gene regulation (7.54%), and specialised metabolism (including plant hormones) (18.86%) (Table S3).

Multi-gene fragments of length >200 kb can be transferred via HGT in grasses (Dunning et al., 2019). Therefore, we checked for co-localising candidates, aiming to find such fragments (Table S3). We found four fragments containing at least three candidate HTGs whose donor clade was the same, three from Chloridoideae and one from Andropogoneae, and six more containing two genes. In all cases, when multiple candidate genes from the same biosynthetic pathway were found, they co-localise, suggesting they are from the same transfer event (Table S3).

We hypothesise that HGTs may be enriched in biosynthetic gene clusters (BGCs). To test this hypothesis, we predicted BGCs in the *Z. bulbosa* genome and found a total of 313 putative BGCs (Dataset S2). Thirteen percent of the candidate HTGs (31 genes including all homologs, out of 237 genes in total) were located within two putative BGCs (benzoxazinoids and strigolactones). To test if the number of candidate HTGs overlapping with BGCs was significantly higher than expected by chance, we performed a permutation test. Under a null model of random genomic distribution, the expected number of overlaps was 5.23 ± 2.28, in contrast with the observed value of 31, around six times higher (Fig. 3). This indicates that HGT-derived genes are non-randomly associated with BGC regions in *Z. bulbosa* (Z-score = 11.32, p-value = 0).

**Figure 3.**
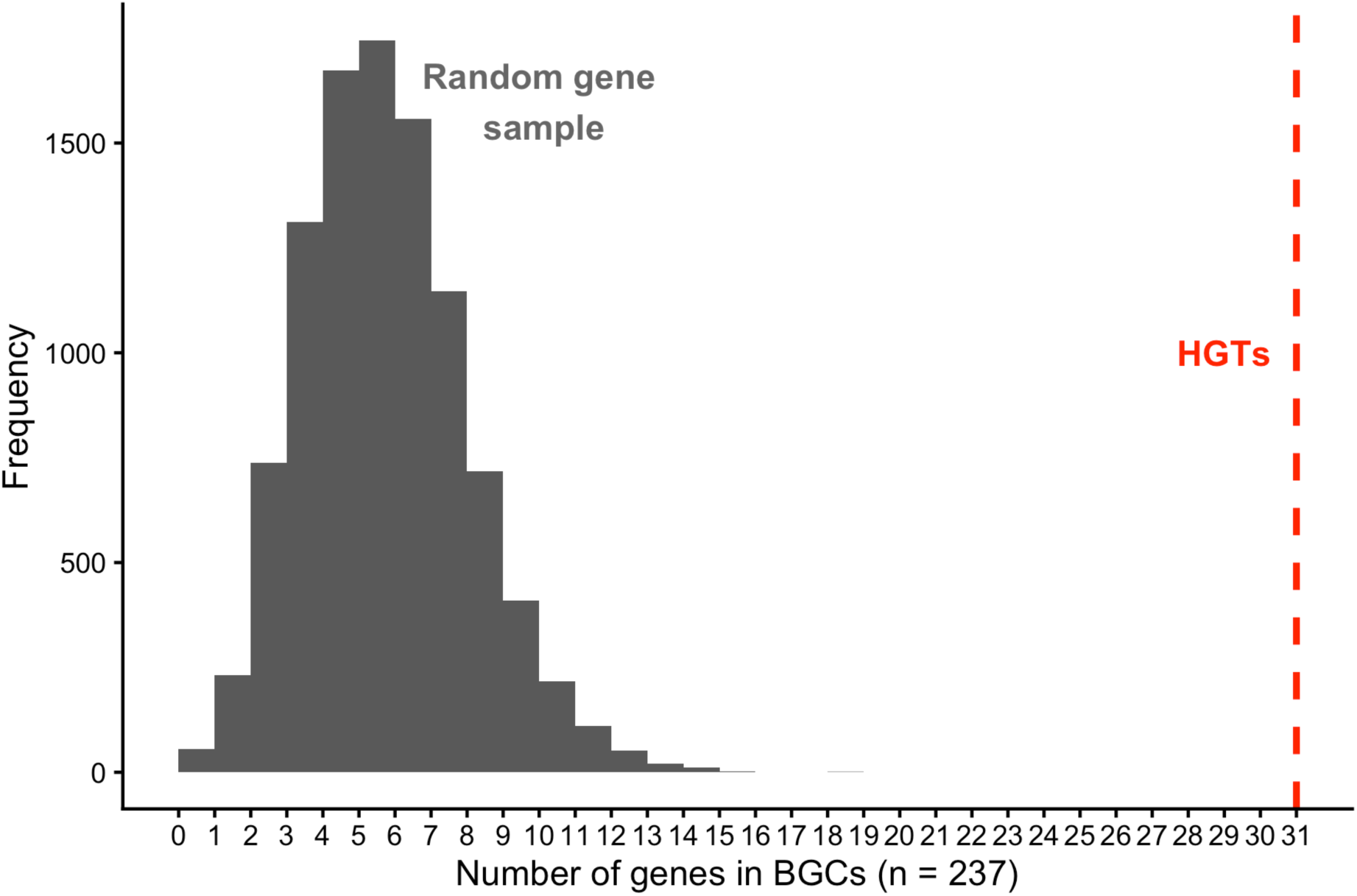
Distribution of overlap between HTG and genes within BGCs in 10,000 random gene sets. The red dotted line is the number of observed overlaps in the *Z. bulbosa* genome.

Among the candidate HTGs within BGCs, we found genes encoding enzymes involved in two biosynthetic pathways. HGT_009 and HGT_042 were annotated as *Bx6* and *Bx7*, from the benzoxazinoid pathway (Jonczyk et al., 2008). HTG_034, which encodes a cytochrome P450 homologous to Arabidopsis MAX1/CYP711A1 (Abe et al., 2014) and HTG_055, which encodes a *S*-adenosyl-methionine-dependent methyltransferase homologous to Arabidopsis carlactonoic acid methyltransferase (CLAMT) (Mashiguchi et al., 2022), are putatively involved in strigolactone biosynthesis. A third candidate, HTG_017, is also part of the *Z. bulbosa* strigolactone BGC, and it is annotated as a cytochrome P450 in the CYP716A subfamily, whose members typically function in triterpenoid biosynthesis (Miettinen et al., 2017). Even though there is currently no evidence for this specific subfamily of cytochrome P450 to be involved in strigolactone biosynthesis, cytochrome-P450 catalysed reactions are core to the pathway diversity (Niu et al., 2025) and the genes are known to be easily recruited from diverse subfamilies (Hansen et al., 2021). Furthermore, given that the gene is located within a previously described BGC, we infer it acts in the same pathway as the other two enzymes.

### Horizontal transfer of the strigolactone biosynthetic gene cluster

A strigolactone BGC has been already described in grasses (C. Li et al., 2024), with genes homologous to the three contiguous candidate HTGs we found in *Z. bulbosa*. Although the bootstrap support was not enough to consider them robust candidates, the fact that they are contiguous in the genome and nested within the same donor clade (Chloridoideae) indicates that very likely they were horizontally transferred but the existence of several paralogs makes the phylogenetic reconstruction more challenging.

We constructed phylogenetic trees for the three genes using an extensive grass genome database (Table S1). With the increased taxon sampling and alternative methods (see Materials and Methods), *Z. bulbosa* strigolactone genes appear robustly nested within the Chloridoideae clade (Fig. 4 and Dataset S1). Despite this, the trees present some incongruencies (e.g. Paspaleae nested within Paniceae instead of sister to Andropogoneae).

**Figure 4.**
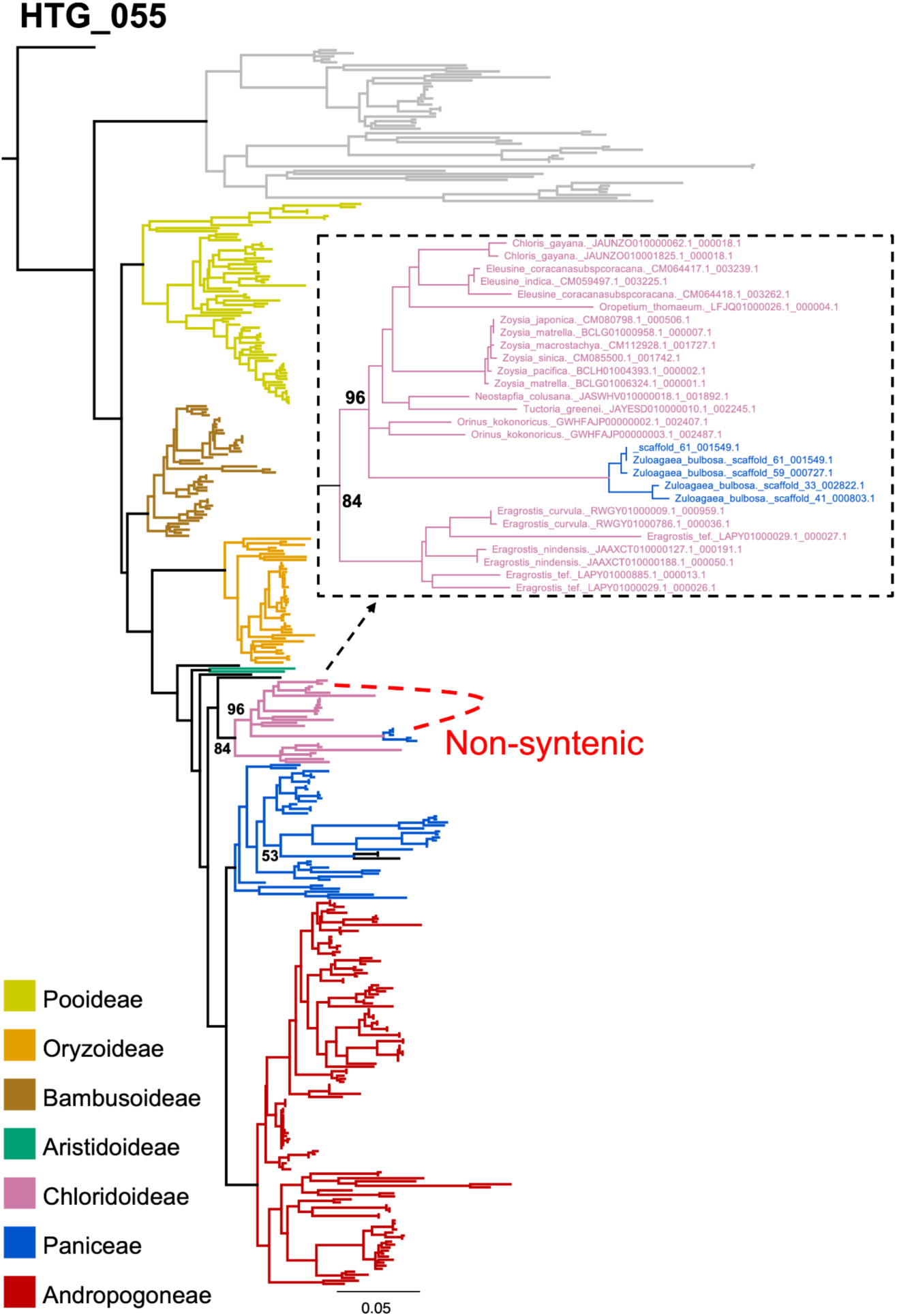
Gene tree with the increased taxon sampling for candidate HTG_055, which is homologous to the strigolactone biosynthetic gene *CLAMT*. The grey clade is an additional paralog.

To further validate that the strigolactone BGC has been horizontally transferred, we investigated the synteny of CLAMT paralogs in *Z. bulbosa*, including vertically inherited and the candidate HTG_055. We confirmed the candidate HTG_055 was not syntenic to the orthologs in *Eleusine coracana*, a Chloridoideae species placed in the donor clade (Fig. 4). In contrast, the vertically inherited *Z. bulbosa* CLAMT are indeed syntenic to the *E. coracana* orthologs. Since genes from Paspaleae species were misplaced in the phylogeny according to what we would expect from the species tree (black clade nested in Paniceae in Fig. 4), we also evaluated their syntenic relationships. Genes from *Paspalum notatum* are syntenic to *E. coracana* orthologs and *Z. bulbosa* vertically inherited copies, indicating that they are likely not HTGs but rather just misplaced in the phylogeny (bootstrap support for that node is only 53).

### A patchy distribution of the benzoxazinoid pathway across grass species

The benzoxazinoid BGC has been thoroughly studied in grasses due to its huge impact in plant defence and resilience (T. Li et al., 2025). Here, we found *Bx6* and *Bx7* orthologs as candidate HTG from Andropogoneae into *Z. bulbosa*. With the aim of gaining understanding in the evolutionary history of this BGC in grasses, we constructed phylogenetic trees for all the genes from the pathway using an extensive grass genome database (Table S1). In most *Bx* genes, we found a topology consistent with the previously documented ancestral HGT from Panicoideae into Pooideae (Fig. 5 and Dataset S1), (Y. Huang et al., 2026; D. Wu, Jiang, et al., 2022). Based on our phylogeny, the ancestral HGT event likely happened before Triticeae and Poeae lineages diverged approximately 45 MYA (Gallaher et al., 2022) (Fig. 5), in contrast with previous reports that placed it after these two lineages split.

**Figure 5.**
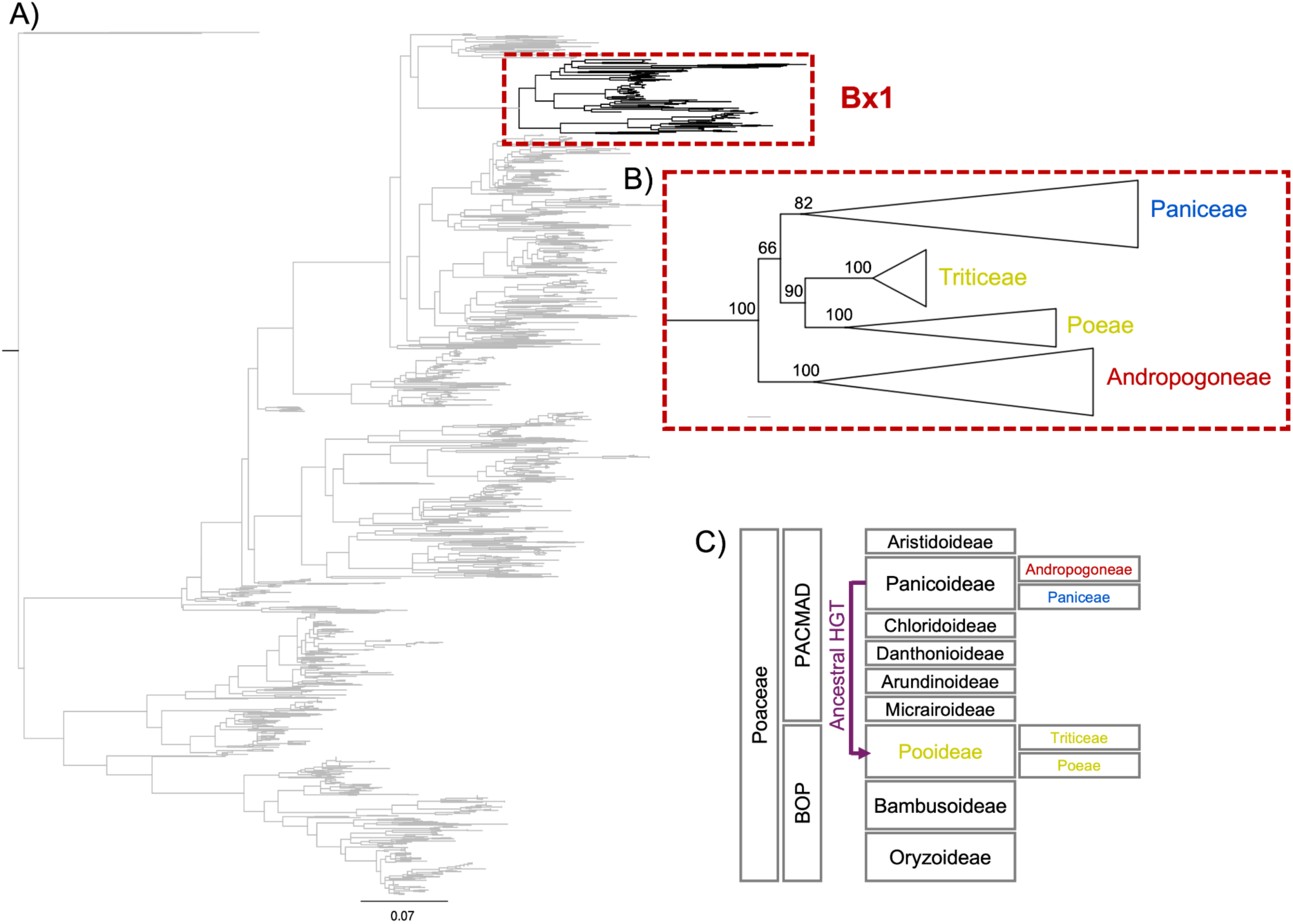
Phylogenetic tree of *Bx1* gene. A) The complete phylogeny in grey includes all homologous sequences in the grass database, used to identify the *Bx1* clade (black). B) Zoomed view into the *Bx1* clade with collapsed clades, showing Triticeae and Poeae lineages nested within Panicoideae. C) Diagram representing the grass phylogeny and the HGT event.

Using the phylogeny, we inferred which genes are present (and how many copies) in grass genomes (Table 2 and Table S4). The Bx pathway presents a patchy distribution, but commonly appears in Andropogoneae, Paniceae, Triticeae, and Poeae subtribes. Yet, in some species from those clades, such as *Sorghum bicolor* from Andropogoneae, the pathway has been completely lost. The broader taxonomic distribution used here allowed us to identify Bx genes present in 19 out of 21 of the included Poeae species (Table S4), with six species having them organised in BGCs (Table S5).

**Table 2.** Analysis of the presence/absence of Bx genes in the 204 grass species with reference genomes identified by phylogenetic methods.

| Taxonomy |  | #<br>gen* | Species containing... |  |  |  | Organised<br>in clusters |
| --- | --- | --- | --- | --- | --- | --- | --- |
|  |  |  | ≤ 2 Bx<br>genes | Partial<br>pathway | Core<br>pathway | Full<br>pathway |  |
| Anomochlooideae |  | 1 | 0 | 0 | 0 | 0 | 0 |
| Pharoideae |  | 1 | 0 | 0 | 0 | 0 | 0 |
| Bambusoideae | Arundinarieae | 5 | 0 | 0 | 0 | 0 | 0 |
|  | Bambuseae | 9 | 1 | 0 | 0 | 0 | 0 |
|  | Olyreae | 3 | 0 | 0 | 0 | 0 | 0 |
| Oryzoideae |  | 19 | 0 | 0 | 0 | 0 | 0 |
| Pooideae | Brachypodieae | 6 | 0 | 0 | 0 | 0 | 0 |
|  | Bromeae | 2 | 0 | 0 | 0 | 0 | 0 |
|  | Poeae | 21 | 6 | 6 | 7 | 0 | 6 |
|  | Stipeae | 1 | 0 | 0 | 0 | 0 | 0 |
|  | Triticeae | 24 | 0 | 0 | 20 | 0 | 0 |
| Aristidoideae |  | 2 | 0 | 0 | 0 | 0 | 0 |
| Arundinoideae |  | 1 | 0 | 0 | 0 | 0 | 0 |
| Chloridoideae |  | 17 | 1 | 0 | 0 | 0 | 0 |
| Panicoideae | Andropogoneae | 54 | 18 | 2 | 3 | 19 | 26 |
|  | Paniceae | 22 | 5 | 4 | 6 | 0 | 10 |
|  | Paspaleae | 3 | 3 | 0 | 0 | 0 | 0 |
|  | Chasmanthieae | 1 | 0 | 0 | 0 | 0 | 0 |

\* Number of genomes per subfamily/tribe.

### Repeated horizontal gene transfer contributed to the assembly of the benzoxazinoid BGC in *Z. bulbosa*

In addition to the ancestral HGT event, our analysis identifies a subsequent, more recent transfer of *Bx6* and *Bx7* from the Rottboelliinae lineage (Andropogoneae) into *Z. bulbosa* (Paniceae, Cenchrineae), which co-exist with native copies (Fig. 6 and 7).

**Figure 6.**
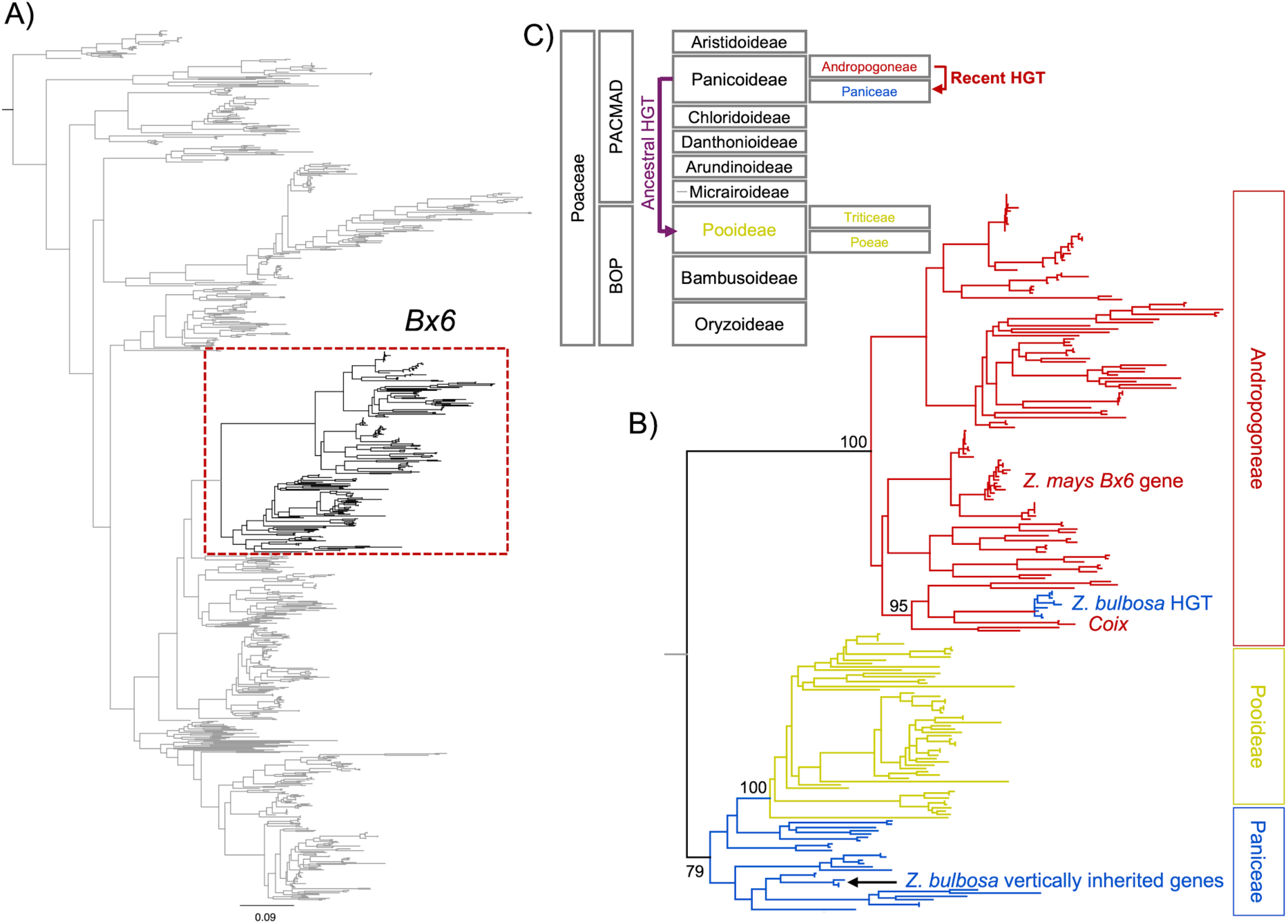
Phylogenetic tree of *Bx6*. A) The complete phylogeny in grey includes all homologous sequences in the grass database, used to identify the *Bx6* clade (black). B) The colored phylogeny zooms into the *Bx6* clade to show the two HGT events, an ancestral one from Paniceae into Pooideae, and a recent one from Andropogoneae into *Z. bulbosa*. C) Diagram representing the grass phylogeny and the HGT events.

**Figure 7.**
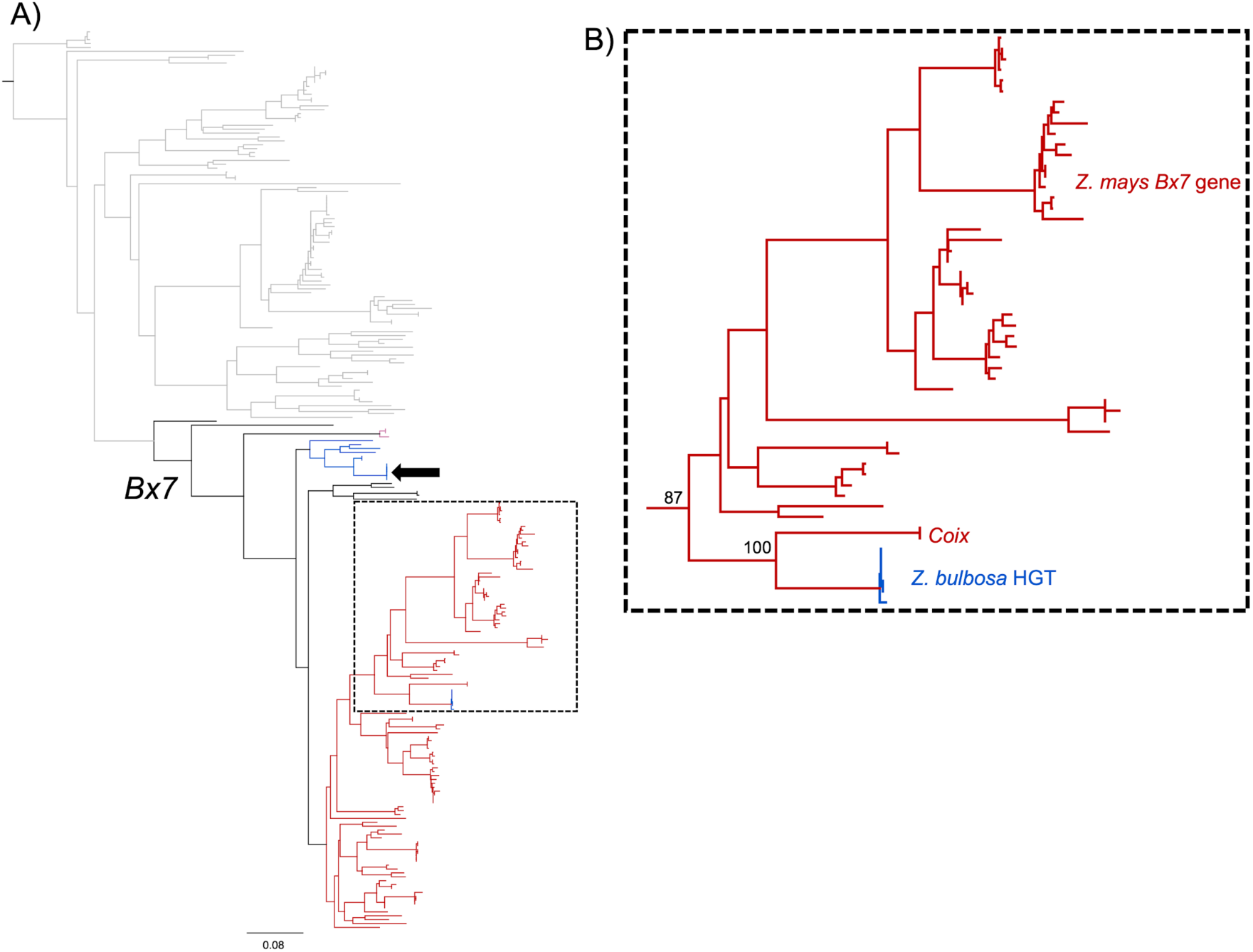
Phylogenetic tree of *Bx7*. A) The complete phylogeny in grey includes all homologous sequences in the grass database, used to identify the *Bx7* clade (black), with the black arrow pointing to *Z. bulbosa* vertically inherited copies. B) The colored phylogeny zooms into the *Bx7* clade to show the recent HGT event from Andropogoneae into *Z. bulbosa*.

In both *Bx6/7* gene trees, *Coix* species appeared as sister of *Z. bulbosa* genes with 100% bootstrap support, suggesting one unique HGT event. The native range of *Coix* spans from South Asia to Australia, making it unlikely to be the actual donor. Assuming that a closely related species without a genome assembly available (and therefore absent from our sampling) could be the donor, we checked the native distribution of other species in the Rottboelliinae clade and found two *Rottboellia* species (*Rottboellia campestris* and *Rottboellia aurita*) that overlap with the native distribution of *Z. bulbosa* in Southern North America, Central America and Northern South America (https://powo.science.kew.org/). *Coix* and *Rottboellia* are estimated to have diverged ∼5 million years ago (MYA) (Welker et al., 2020), so if an American *Rottboellia* species is the donor, the HGT event would be very recent in contrast to the ancestral HGT from Panicoideae into Pooideae (Y. Huang et al., 2026; D. Wu, Jiang, et al., 2022). The HTGs are absent from closely related Cenchrinae species, such as *Setaria italica* and *Setaria viridis*, which do have a native *Bx6* copy, also pointing to a recent transfer.

### The genomic structure of benzoxazinoids biosynthetic gene cluster is conserved in divergent grasses

The benzoxazinoid BGC was identified in maize, and it was the first one to be characterised in plants (Frey et al., 1997). Since then, *Bx* orthologs have been found to be often localised on the same chromosome/scaffold, suggesting that the pathway is mostly organised as a BGC in grasses (D. Wu, Jiang, et al., 2022). The core pathway (Bx1-5 and Bx8, which are the genes clustered in maize) was contained in the same chromosome in six Poeae species, ten Paniceae species and 26 Andropogoneae species (Table 2 and Table S5), showing a remarkable conservation of the genomic architecture of the pathway. The exception is found in Triticeae species, whose *Bx* genes were not clustered, despite containing the full functional pathway, as previously described (Sue et al., 2011; D. Wu, Jiang, et al., 2022). To compare the BGC structure for *Bx* genes across different grasses, we predicted and visualised BGCs in three representative species: *Z. bulbosa* (Paniceae), *Vossia cuspidata* (Andropogoneae), and *Poa pratensis* (Poeae) (Fig. 8A).

**Figure 8.**
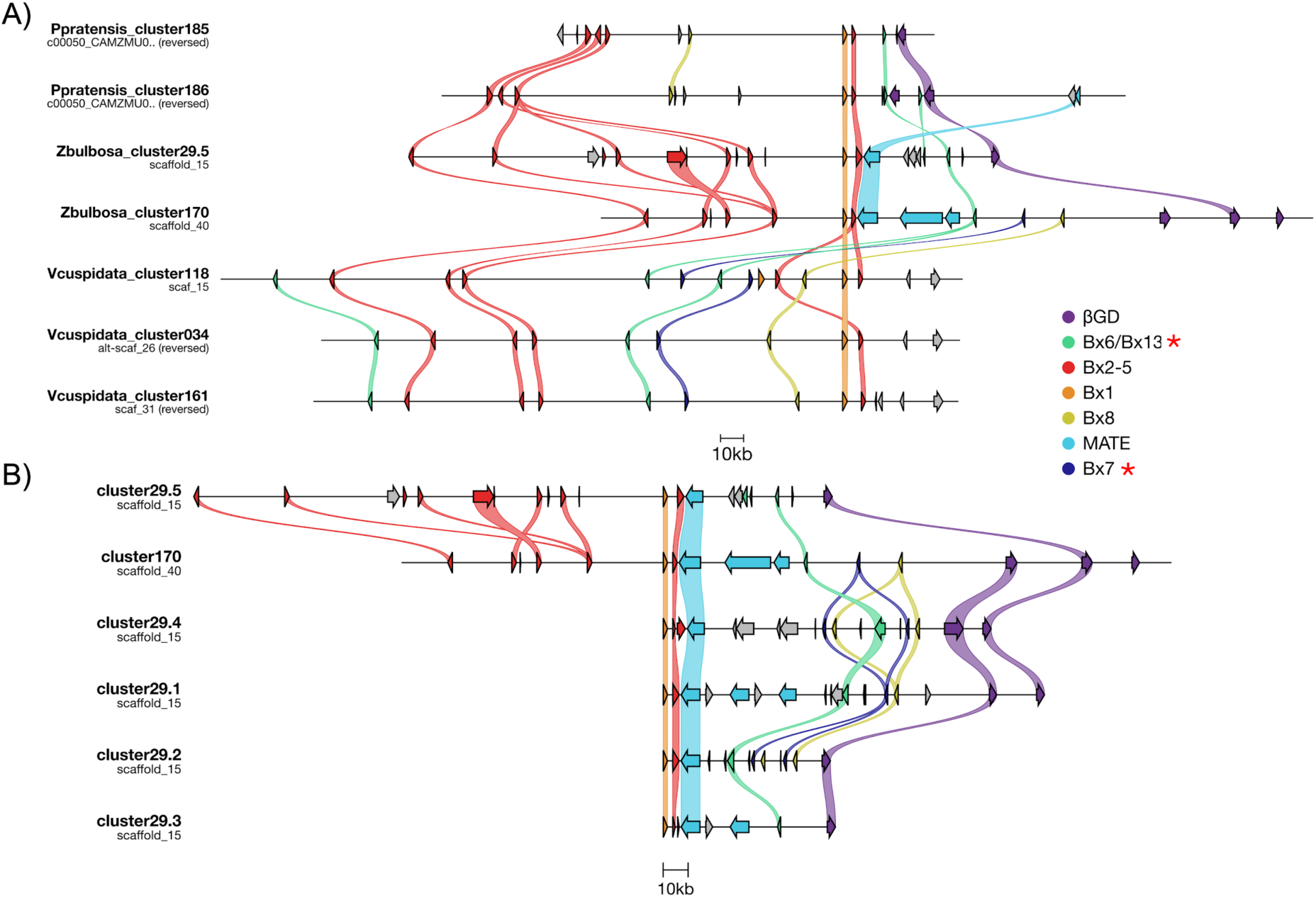
Genomic structure of *Bx* gene cluster in grass species. A) All clusters in *P. pratensis, Z. bulbosa*, and *V. cuspidata* are represented. For cluster 29 from *Z. bulbosa*, only one tandem repeat is shown. B) *Z. bulbosa* clusters, with cluster 29 split into its tandem repeats (subclusters). The red asterisks indicate the gene classes that are HTGs in *Z. bulbosa*. The domains predicted by PlantiSmash (Table S6) were used to infer homology with Bx genes and are represented in different colors. In both figures, *Bx1* homologs are used as an anchor to display the BGC.

In *Z. bulbosa*, two Bx BGCs were predicted, both with copies of the candidate HTGs. Most of the genes contained in the BGC were vertically inherited (Fig. 5 and Dataset S1), except horizontally transferred *Bx6* and *Bx7*. Surprisingly, their corresponding native, vertically inherited copies were dispersed on other chromosomes. In addition to the previously characterised genes, the BGC contained eight multidrug and toxic compound extrusion (MATE) transporters and seven glycosyl hydrolases or **β**-glucosidases (*βGD*). The genomic organisation of cluster 029 suggests several tandem duplications of the original cluster, leading to a ‘super-cluster’ that spans ∼815 kb (Table S6 and Fig. 8B), followed by extensive genomic rearrangements including gene loss of some of the genes in specific subclusters. A more compact, mostly single-copy *Bx* BGC (cluster 170) is present in a different genomic location (Table S6 and Fig. 8B), which also contains the horizontally acquired copies of *Bx6* and *Bx7*.

The genome of *P. pratensis*, from the Poeae lineage within Pooideae, contains two Bx BGCs on the same scaffold (CAMZMU010000065.1), separated by 163 kb (Table S6). Both BGCs contain nearly the complete pathway, with the exception of *Bx7* (Fig. 8A). Interestingly, glycosyl hydrolases and one MATE transporter co-localise as well with the known *Bx* genes. In contrast, the *Bx* clusters found in the genome of *V. cuspidata*, from the subtribe Rhytachninae within Andropogoneae, were similar to the one described in maize and did not contain glycosyl hydrolases nor MATE transporters, although *Bx6* and *Bx7* were still embedded in the BGCs (Fig. 8A and Table S6).

## Discussion

### Horizontal gene transfer between *Z. bulbosa* and maize is reciprocal

Grass-to-grass HGT is extensive (Hibdige et al., 2021; Y. Huang et al., 2026) however the dynamics of this gene flow are not well-known, in part due to the lack of a proven mechanism (Pereira et al., 2022). The rates of acting as donor and/or recipient species vary across the Poaceae family (Hibdige et al., 2021; Y. Huang et al., 2026). These rates seem to be correlated, meaning that certain taxa, such as the Paniceae and Andropogoneae tribes, are more likely to be both donors and recipients (Hibdige et al., 2021; Y. Huang et al., 2026). In addition, phylogenetic distance and geographical co-occurrence seem to be correlated with the number of transfers (Hibdige et al., 2021), yet the factors contributing to this variation are not well understood.

Here we present the first case of bidirectional HGT between two distant grass species, *Zea mays* and *Zuloagaea bulbosa* (Fig. 1 and Fig. 2). This system is especially relevant because maize is one of the most important cereal crops worldwide while *Z. bulbosa* is sometimes considered as a weed in or near agricultural land. The interchange of genetic material between crops and weeds may have profound implications in how we manage agricultural systems and even in considering the likelihood of gene escapes from transgenic crops into weeds (Pilson & Prendeville, 2004).

### Horizontal gene transfer contributes to the evolution of biosynthetic gene clusters

Our results in *Z. bulbosa* suggest that there is a correlation between HGT and gene clustering (Fig. 3). This association between gene clusters and HGT is well established in bacteria (Dilthey & Lercher, 2015; Homma et al., 2007) and fungi (Rokas et al., 2020; Wisecaver & Rokas, 2015). In fungi, which are eukaryotes and display a rich and complex specialised metabolism like plants, genes in metabolic gene clusters (biosynthetic and other metabolic pathways such as primary metabolism) were shown to be horizontally transferred 1.66-fold more often than their non-clustered counterparts (Wisecaver et al., 2014; Wisecaver & Rokas, 2015). When analysing “gene neighbourhoods” independently of the function of the underlying genes, their evolution was mostly by vertical inheritance, except for metabolic gene clusters, which were significantly more affected by HGT (Marcet-Houben & Gabaldón, 2019). Here we showed a similar result in a grass genome, although the low frequency of HGT events in plants is a limitation. Future studies need to extend such analyses to a broader taxonomic representation of plants to confirm whether the correlation between HGT and BGCs extends beyond *Z. bulbosa*, especially given that both these processes are potential drivers of metabolic innovation.

The enrichment in BGC of HTGs may be due to the immediate and strong adaptive benefit(s) they could have post-transfer. Maintaining polygenic traits through tight linkage might be particularly useful when the trait confers a locally adaptive advantage (Smit & Lichman, 2022). Presumably, BGCs in the donor species are already under strong positive selection, and sympatric grasses face similar biotic and abiotic stresses. Since a complete ‘functional unit’ - full biosynthetic pathway - is transferred in one event, likely with its own cis-regulatory elements (Zheng et al., 2015), the foreign DNA is ready to be used (Collins et al., 2026) and the gained phenotypic trait may immediately contribute to fitness gains. Therefore, perhaps HGTs do not happen more frequently in BGCs, but they are just retained more than other neutral or deleterious transfers due to their modular structure (Van Etten & Johnson, 2026).

An alternative explanation could be that the molecular mechanism leading to DNA movement and integration in a different genomic location, which is required for BGC construction, could be similar to the processes that lead to genomic integration required for HGT. Transposable elements are thought to mediate the gene movement required for BGC assembly, either through ectopic recombination or active transposition (Smit & Lichman, 2022). The horizontal transfer of transposable elements is pervasive, being detected more frequently and broadly in terms of taxonomy than protein-coding genes (Aubin et al., 2021; El Baidouri et al., 2014; Park et al., 2021). In this study we found two transposable element-like sequences horizontally transferred, despite not targeting these in our analyses. So, if activation and mobilisation of transposable elements are involved in both HGT and BGC assembly, the temporal and spatial context might be the same and lead to the overlaps we observe.

Beyond the enrichment pattern, we showed two specific examples of BGC evolution shaped by HGT in *Z. bulbosa*. The transfer of *Bx6* and *Bx7*, which belong to the benzoxazinoid pathway, into an already established BGC, illustrates the fact that HGT can not only distribute existing BGCs (Y. Huang et al., 2026; D. Wu, Hu, et al., 2022; D. Wu, Jiang, et al., 2022) but also actively contribute to BGC evolution, potentially modulating the efficiency and/or the final products of the pathway. We also found a horizontally transferred fragment containing three non-homologous genes that were part of a strigolactone BGC identified in *Z. bulbosa* and in a donor proxy, the Chloridoideae species *Eleusine coracana* (Fig. 4 and Table S6). This BGC has been validated in rice (C. Li et al., 2024), and it seems to have homologous BGCs in other grasses (Niu et al., 2025). Strigolactones are a group of plant signalling molecules of terpenoid origin with multiple functions in plant development and environmental interactions (Mashiguchi et al., 2021). The ‘backbone’ of strigolactone biosynthesis is well known, but the pathway diversifies downstream to produce chemically diverse compounds through decoration of the intermediate molecule, carlactone (Vinde et al., 2022). The dispersal of the BGC from Chloridoideae as a functional unit through HGT could directly diversify the metabolic array of *Z. bulbosa*, offering new avenues for adaptation.

### Biosynthetic gene clusters are highly dynamic and evolve quickly

Gene movement via genomic rearrangement is essential for the emergence of BGCs. Indeed BGCs are usually located in dynamic chromosomal regions rich in transposable elements (Field et al., 2011). Common genomic rearrangements, such as insertions, inversions and duplications, are frequently observed in BGCs across the plant kingdom (Q. Li et al., 2020; Liu et al., 2023).

Here we revealed the diversity in the genomic architecture of a paradigmatic BGC in grasses (Fig. 8 and 9). Multiple steps of re-shuffling, including not only the movement of native DNA but also the horizontal transfer of key genes from other distantly related grass species, are predicted to have contributed to the pathway assembly and its dispersal to other species through HGT. It is also predicted that extensive gene losses also took place over evolutionary time, eradicating the pathway from several grass clades, and even specific species within clades that usually carry the pathway. The cluster architecture has been maintained over ∼45 MYA in divergent species from Paniceae, Andropogoneae and Poeae, whilst in Triticeae the pathway is conserved but the genes have been dispersed across several chromosomes.

**Figure 9.**
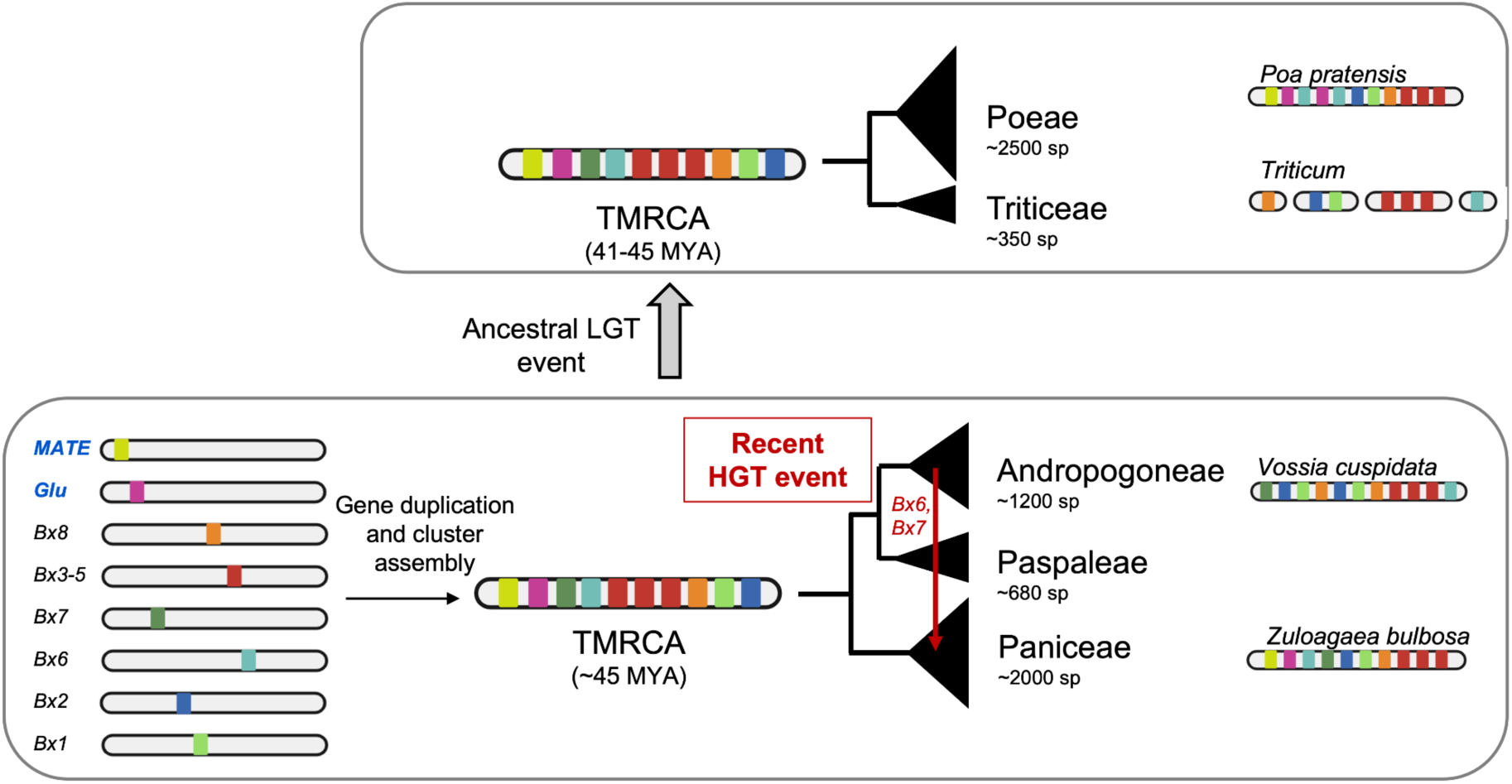
Hypothesis of the evolutionary history of the benzoxazinoid biosynthetic gene cluster (modified from (D. Wu, Jiang, et al., 2022)). An extended cluster, including *βGD* and MATE transporters, was assembled in the most recent common ancestor (TMRCA) of Andropogoneae, Paspaleae and Paniceae and transferred horizontally into the most recent common ancestor of Poeae and Triticeae around 45 MYA. Another, more recent HGT event from Andropogoneae moved horizontally acquired copies of *Bx6* and *Bx7* into the *Z. bulbosa* BGC. Despite pervasive gene losses across the grass phylogeny, the BGC has been conserved over millions of years in certain species, whilst it has beenre-shuffled in Triticeae species.

Furthermore, tandem duplications of both individual genes and full clusters were found, with an especially remarkable example in *Z. bulbosa*, where five divergent copies of the cluster were found in tandem (Fig. 8B). The rapid evolution of BGCs, which act as functional units usually leading to an adaptive phenotype, may be due to the co-localisation itself. High rates of gene duplication generate the substrate that allows for neo and subfunctionalisation of biosynthetic genes and pathways. For example, the steroid triterpenoids withanolides are synthesised in aerial and below-ground tissues by two homologous but distinct BGCs located next to each other (Priego-Cubero et al., 2025). Although further experimental validation is needed to see if the duplicated copies are subfunctionalised, we showed an analogous genomic architecture for the *Bx* cluster in *P. pratensis* and *Z. bulbosa*.

### Transporters and β-glucosidases belong to the benzoxazinoid biosynthetic cluster in divergent grasses

In addition to the known *Bx* genes, we found two more gene families, *βGD* and MATE transporters co-localising with the *Bx* BGC. *βGD* are functionally related with the Bx pathway (Czjzek et al., 2000), but MATE transporters for benzoxazinoids have not been found before. These genes, although they are not strictly involved in the biosynthesis of benzoxazinoids, may modulate the spatial distribution of the active, toxic compounds, and therefore they are likely to be relevant for plant protection. They are located within the *Bx* BGC in several, distant species from the Poeae and Paniceae tribes, indicating that they were initially clustered and presumably dispersed later in the Andropogoneae (Fig. 8 and 9).

*βGD* (glycosyl hydrolases) remove the glucose from the conjugate DIMBOA-Glc, liberating the active compound, DIMBOA (Czjzek et al., 2000). Several paralogs encoding these enzymes in maize have been characterised and are located on chromosomes 3 and 10, outside of the *Bx* BGC (Gómez-Anduro et al., 2011). However, in other grass genomes, such as *Z. bulbosa* and *P. pratensis*, these genes co-localise with the rest of the biosynthetic pathway. When inspecting the phylogenetic trees for glycosyl hydrolases, we found that the topology was similar to those found in other Bx genes (Fig. S3), indicating that they were originally contained in the BGC and later dispersed in Andropogoneae species, as it happened for *Bx6* and *Bx7*. Only the **β**-glucosidases from certain species (such as *Z. bulbosa*, *Dichantelium oligosantes*, *Echinochloa colona*, and *Poa* species) co-localise with the BGC, whilst others, including in maize, have *Glu* genes dispersed across the genome.

MATE transporters commonly move specialised metabolites across membranes, allowing for the control of metabolite accumulation in a tissue and in an organelle-specific manner in plants (Nour-Eldin & Halkier, 2013). Other BGCs, such as the dhurrin cluster in *Sorghum* (Darbani et al., 2016), the cucurbitacin cluster in cucumber (Ma et al., 2023), and the nicotine cluster in tobacco (Schwabe et al., 2026) contain MATE transporters, but no transporters have been described yet for benzoxazinoid compounds (Niculaes et al., 2018). We did find MATE transporters within *Z. bulbosa* and *P. pratensis* BGCs, indicating that they are likely involved in the extrusion of toxic compounds of the Bx pathway (Table S6). The paralog identified in the *P. pratensis* BGC shows a topology compatible with the hypothesis of ancestral HGT followed by multiple gene losses, since Poeae and Paniceae species appear as monophyletic with 100% bootstrap support, but Triticeae and Andropogoneae lack this paralog (Fig. S4). These paralogs are located on the same chromosomes/scaffolds as the *Bx* genes in Poeae and Paniceae species (Table S5). Surprisingly, the paralog clustered in *Z. bulbosa* BGCs is quite distant in the phylogeny (Fig. S4), pointing to an independent recruitment.

## Conclusion

Horizontal gene transfer allows for a limited gene flow across species boundaries, widening the standing genetic variation that selection can act upon. We here show that this transfer can be bidirectional, with the same pair of species acting as both donor and recipient. Although most HTGs are neutral and purged over time, a small proportion may be adaptive. We found HTGs from *Z. bulbosa* predominantly located within BGCs, probably due to their modular structure allowing for an immediate adaptive benefit which undergoes strong positive selection. The dynamic nature of BGCs, which commonly undergo a myriad of genomic rearrangements including horizontal gene transfer, promotes the rapid and diversifying evolution of plant specialised metabolism.

## Materials and methods

### Plant material

Seeds from *Zuloagaea bulbosa* (Kunth) Bess accession PI422481 from USDA Germplasm Resources Information Network (GRIN) were germinated in a Petri dish with damp filter paper for ∼10 days in a Sanyo MLR-352H growth chamber (12 h light and 20°C, 12 h dark and 15°C and 70% humidity). After radicle and shoot emergence, the seedling was transferred to a small pot filled with F2S standard soil and a lid was used to maintain high humidity for acclimation. When the plant was more established, after about 2-4 weeks, it was transferred to a 2 L pot with a 4:1 ratio of F2S standard soil:perlite (East Riding Horticulture Ltd) and 8 gof slow-release fertiliser was added (East Riding Horticulture Ltd). The plant grew in a glasshouse at the Arthur Willis Environment Centre, University of Sheffield (UK) under semi-controlled conditions (12 h daylight, 25/20 °C day/night temperature).

Young leaves were collected from one individual *Z. bulbosa* plant for high molecular weight (HMW) DNA extraction for HiFi and for Hi-C sequencing. Tissue was placed in 50 mL Falcon tubes and immediately frozen in liquid nitrogen and then kept at -80°C until used.

### DNA extraction and sequencing

Frozen leaves were ground to a fine powder in liquid nitrogen with a pestle and mortar. DNA was extracted from ∼500 mg of ground tissue using the Nucleobond® HMW DNA kit (Macherey-Nagel) following manufacturer’s instructions. The DNA pellet was left 3 days at room temperature to resuspend, and then kept at 4 °C. DNA concentration, purity, and integrity was verified with Nanodrop (Thermo Fisher Scientific), Qubit (Thermo Fisher Scientific), and Femtopulse (Agilent). Samples were sent for library prep and HiFi sequencing on a Sequel II PacBio instrument (three flow cells) at the Centre for Genomic Research at the University of Liverpool (United Kingdom).

For Hi-C sequencing, we sent the collected frozen leaves (∼2 g of fresh material) to BGI Genomics (China). They performed the cross-linking with formaldehyde, digested the DNA with a restriction enzyme and incorporated biotinylated residues to the 5’ overhangs. The blunt-ends were ligated to obtain circular molecules and these were purified. The target circular DNA was sheared and captured through biotin-streptavidin‒mediated pulldown. Hi-C libraries were generated and sequenced on a DNBSEQ sequencing platform (PE150 reads). The raw reads were filtered by BGI Genomics with an in-house developed software called SOAPnuke (Y. Chen et al., 2018). To retain only high quality reads, reads with length ≤150 (- -minReadLen 150), with polyX ≥ 50 bp (--polyX 50), N content ≥ 1% (-n 0.01), ≥25% of the sequence corresponds to adapters (--adaMR 0.25), ≥30% of the bases had a quality lower than 20 (-l 20) or the quality was ≤ Phred+33 (-q 0.3), were discarded.

### Genome size estimation

The 1C genome size for each species was estimated by flow cytometry using the one-step protocol (Doležel et al., 2007) with minor modifications (Clark et al., 2016). *Petroselinum crispum* ‘Champion Moss Curled’ (2C = 4.50 pg) (Obermayer et al., 2002) was used as the calibration standard and nuclei were isolated in the General Purpose Buffer (Loureiro et al., 2007), supplemented with 3% PVP and 0.08% (v/v) beta-mercaptoethanol.

### Genome assembly and annotation

We used HiFi data to obtain a k-mer based estimate of genome size and ploidy composition using Genomescope v2.0 and Smudgeplot v0.4.0dev (k=21) (Ranallo-Benavidez et al., 2020). To assemble a draft genome for *Z. bulbosa*, we first used both the HiFi and Hi-C data as inputs for hifiasm v0.19.9-r616 with the number of haplotypes set to 2 (Cheng et al., 2021). Due to the complex polyploid nature of *Z. bulbosa* (as triploids, hexaploids and octoploids have been reported based on chromosome counts, (Rice et al., 2015)) we used the primary unitigs (haplotype-resolved processed unitig graph without small bubbles) as a reference to align Hi-C data. We performed an additional cleaning step of Hi-C short-reads with fastp v1.0.1 using default parameters (S. Chen, 2023). For scaffolding we first mapped the cleaned Hi-C reads to the primary unitigs with bwa v0.7.19-r1273 (H. Li & Durbin, 2010), filtering for multi and low quality mapped reads (MAPQ ≥30). We then used the YAHS v1.2.2 pipeline, skipping the initial assembly error correction step (--no-contig-ec flag) (Zhou et al., 2023). We visualised the contact map using juicer_tools v1.9.9 and Juicebox v2.15 (Dudchenko et al., 2018; Durand et al., 2016), which are part of HapHiC pipeline. Assembly quality control was performed using Quast v5.2.0 (Mikheenko et al., 2018) for contiguity and BUSCO v6.0.0 (Simão et al., 2015) with poales_odb10 dataset for completeness. The genome assembly was annotated using Helixer v0.3.6 (Holst et al., 2023) with the land_plant database and a batch size of 512, and coding and protein sequences extracted with gffread v0.12.7 (Pertea & Pertea, 2020). Protein functional annotation was performed on InterProScan v5.73-104.0 (Jones et al., 2014).

To mark scaffolds derived from organelle DNA, regions from the *Z. bulbosa* assembly with high homology to grass chloroplast DNA (NCBI Reference Sequences NC_036384.1 - *Cenchrus purpureus*, NC_028075.1 - *Setaria viridis*, NC_030067.1 - *Urochloa brizantha*, and NC_001666.2 - *Zea mays*) and mitochondrial DNA (NCBI Reference Sequences AP012527.1 - *Oryza rufipogon*, and NC_086594.1 - *Paspalum vaginatum*) were identified with blastn (Ye et al., 2006). We also mapped the HiFi reads to the assembly using minimap2 v2.30-r1287 (H. Li, 2018) and computed the coverage using samtools v1.21 coverage (Danecek et al., 2021). Scaffolds that had ≥80% of their sequence matching to chloroplast DNA with a pairwise identity of ≥90% were marked as chloroplast scaffolds. Scaffolds that had ≥25% of their sequence matching to mitochondrial DNA with pairwise identity of ≥90%, and ≥30x average coverage were marked as mitochondrial scaffolds. Laxer parameters were used for the mitochondrial genome compared to the chloroplast genome because of the high complexity and variability of plant mitochondrial genomes, which frequently diverge between species and contain DNA originating from other cellular compartments like the chloroplast and the nucleus (Z. Wu, Liao, et al., 2022).

### Synteny analyses

Macrosynteny between the *Z. bulbosa* genome and the *Setaria viridis* reference genome was assessed using whole-genome alignment followed by dotplot visualization. Pairwise sequence alignments were generated using minimap2 v2.30-r1287 (H. Li, 2018) and the output results were plotted as dotplots using the R script dotPlotly (https://github.com/tpoorten/dotPlotly, accessed 05/2026).

Collinear blocks between *Z. bulbosa*, *Paspalum notatum* and *Eleusine coracana* were inferred. In brief, the amino acid sequences of all predicted genes for each pair of species were blasted against each other using default parameters, and this was used as an input to calculate collinear blocks with MCScanX (Wang et al., 2012).

### Horizontal gene transfer identification

To identify candidate horizontally transferred genes (HTG) in the *Z. bulbosa* genome, we used a previously developed bioinformatic pipeline (Dunning et al., 2019; Hibdige et al., 2021; Raimondeau et al., 2023) available from GitHub (https://github.com/Sheffield-Plant-Evolutionary-Genomics/current-LGT-pipeline). In brief, we first use blastn with default parameters to identify *Z. bulbosa* coding sequences with a non-Cenchrinae top-hit match (>300 bp) against a database of 67 genomes (Table S1). For this reduced set of genes, the blast matches were aligned to the query from *Z. bulbosa* using the ‘add_fragments’ flag in MAFFT v7.453 (Katoh & Standley, 2013). If the blastn match for a species was fragmented, the different fragments were joined into a single sequence after they had been aligned using a custom perl script. We then built a maximum likelihood phylogenetic tree using PhyML v.21031022 for each candidate, with the best substitution model identified using Smart Model Selection SMS v.1.8.1 (Lefort et al., 2017) and midpoint rooted the resulting topologies with Phytools in R. The rooted trees were used to check if the sequence was nested outside of Cenchrinae in another well-supported clade of grasses with at least 50% bootstrap support. Finally, for candidates that pass this filter, we built a more densely sampled phylogenetic tree using publicly available transcriptomes (Table S1). We then manually inspected the multiple sequence alignments and phylogenetic trees to verify the gene was horizontally transferred.

For this, all alignment and tree manipulations were done in Geneious® 2025.1.2. First, we joined discontinuous blast matches to the same transcript into one unique sequence. Then, the alignments were re-aligned as codons using MAFFT v7.453 (Katoh & Standley, 2013) and manually trimmed with a codon-preserving method to remove poorly aligned regions. We inferred phylogenies using PhyML v.21031022, with the best substitution model identified using Smart Model Selection SMS v.1.8.1 (Lefort et al., 2017) and manually inspected them. We rooted them using outgroups (the most divergent species from *Z. bulbosa* present in the tree) and we classified them as robust candidate HTGs when (1) more than three species were present both within and outside the donor clade, (2) the candidate HTG was clearly nested with ≥70 bootstrap support with the denser sampling, and (3) there were no evident paralogy problems within the tree.

To check if candidate HTGs were likely part of a horizontally transferred fragment, we searched for candidates that were co-localising (<200 kb from each other) and whose clade donor was the same. When multiple copies of a candidate HTG were present, we collapsed these copies into one group of candidate HTGs (referred as HTG_000), since they most likely represent genes duplicated post-transfer or homeolog sequences due to polyploidisation instead of independent HGT events. In order to be considered a multi-gene fragment, candidate HTGs must be non-homologous (different HTG groups).

The candidate HTGs were used as queries to perform blastx against the Swissprot database (downloaded 19-11-2025). For each sequence, we only kept the best match (highest bitscore). The description of the subject sequence was used as a tentative functional annotation. A literature review was performed for each candidate to assess their function.

### Enrichment of horizontally transferred genes in biosynthetic gene clusters

To predict biosynthetic gene clusters (BGC) in grass genomes, we used PlantiSmash 2.0 software (Del Pup et al., 2025) in the online server (https://plantismash.bioinformatics.nl/). Genome assembly and annotation files were used as inputs for target species, and the PlantiSmash 2.0 pipeline was run using default parameters. The outputs were contrasted with the phylogenetic trees to verify that the genes found in clusters were orthologous to the characterised genes in biosynthetic pathways. The GenBank files generated by PlantiSmash were used to visualise BGCs using clinker v0.0.31 (Gilchrist & Chooi, 2021). Only the best links were shown and identity threshold was 0.3

To test whether horizontally transferred genes in the *Z. bulbosa* genome are enriched for BGCs we used a permutation test. The null hypothesis was that candidate HTGs were randomly distributed across the genome. We generated 10,000 random gene sets of equal size to the observed number of candidate HTGs (n = 237 genes). This number includes all copies for candidate HTGs, excluding HTG_003 and HTG_044, which were transposable elements, and HTG_026, which was a plastid sequence (Table S2). The background set of genes was all nuclear genes (organelle-derived contigs were excluded).

All genes were classified as contained or not contained within BGC by using the predicted clusters (PlantiSmash 2.0 results for *Z. bulbosa* genome). We then recorded the observed number of genes within BGC and the simulated value in the 10,000 permutations. The empirical p-value was calculated as the proportion of permutations yielding a value equal to or greater than the observed overlap. The standardized effect size (Z-score) was calculated as the observed value minus the average of the simulated values, divided by the standard deviation of the simulated values. The distribution was plotted in R using ggplot (Wickham, 2009).

### Presence of benzoxazinoid biosynthetic genes in grasses

We built phylogenetic trees for each benzoxazinoid (Bx) biosynthetic gene to track their presence/absence in 204 grass species with reference genomes (Table S1). To ensure the gene annotation was reproducible across genomes, we re-annotated all the available genomes with Helixer v0.3.6 (Holst et al., 2023). Then, we used all the maize *Bx* coding sequences as queries (*Bx1-Bx14*) to blast against all grasses, using coding sequence fasta files as nucleotide databases. We used relaxed parameters to get relatively distant paralogs (blastn -word_size 9 -gapopen 3 -penalty -2 -gapextend 2), selecting only matches with a minimum length of 300 bp. We aligned these blast matches sequentially (--addfragments) to the initial query using MAFFT v7.453 (Katoh & Standley, 2013) with the L-INS-i algorithm (--localpair --maxiterate 1000). If the blastn match for a species was fragmented, the different fragments were joined into a single sequence after they had been aligned using a custom perl script. Finally, maximum-likelihood phylogenetic trees were inferred using IQ-TREE v2.4.0 (Minh et al., 2020), with the best-fitting substitution model selected using ModelFinder, and branch support assessed with 1,000 ultrafast bootstrap replicates. These trees, which contained a mean of 1,157 tips, were manually inspected in Geneious® 2025.1.2 to find the clade containing the maize Bx gene and observe if its topology was inconsistent with the species tree, therefore presenting signals of ancestral or recent HGT.

To infer the presence/absence and copy number of Bx genes across the grass family, we extracted the tip labels for the Bx clade. Then, we counted the number of copies of each gene in each species and checked whether genes were co-locating in the same chromosome using the packages dplyr (Wickham, François, et al., 2026) and tidyr (Wickham, Vaughan, et al., 2026) in R. The Bx genes were considered to be a BGC if the number of non-homologous Bx genes co-localising in the same chromosome was ≥ 6, since this is the number of genes that maize Bx cluster contains (*Bx1-5* and *Bx8*).

## Acknowledgements

We thank the staff of the Arthur Willis Environment Centre and all members of the Dunning lab for their help with plant growth and maintenance. We thank Lucy Knowles for her support in DNA quality assessment. Sequencing data generation was carried out by the Centre for Genomic Research, which is based at the University of Liverpool. This work was funded by the Natural Environment Research Council (grant no.: NE/V000012/1 and UKRI2660) and NERC environmental omics facilities (grant no.: NEOF1545). LTD is funded by a NERC fellowship (grant no.: NE/T011025/1).

## Supplementary material

**Supplementary Figure S1**. Estimation of genome size and ploidy from k-mer distribution.

**Supplementary Figure S2.** Synteny plot between *Setaria viridis* and *Zuloagaea bulbosa*.

**Supplementary Figure S3.** Phylogenetic tree for **β**-glucosidase gene family in grasses.

**Supplementary Figure S4.** Phylogenetic tree for MATE transporter gene family in grasses.

**Supplementary Table S1.** Genome databases used for the different analyses, including taxonomy and source of the data.

**Supplementary Table S2.** List of candidate HTGs and corresponding details about quality filtering.

**Supplementary Table S3.** List of all individual homolog genes belonging to each HTG group.

**Supplementary Table S4.** Presence/absence and gene copy number for all known genes from the benzoxazinoid pathway in the grass database.

**Supplementary Table S5.** Number of *Bx* genes co-locating in the same chromosome/scaffold in grass species.

**Supplementary Table S6.** Benzoxazinoid and strigolactone BGC predicted by PlantiSmash in selected grass species.

## Data availability

The *Z. bulbosa* raw data, genomes and annotation files are available in NCBI (Bioproject PRJNAxxxxx). InterProScan functional annotation, tree files and BGC predictions from PlantiSmash (Datasets S1, S2 and S3) are available on dryad (doi: xxxxxxx). All code generated for this study is available in GitHub (https://github.com/Sheffield-Plant-Evolutionary-Genomics).

## Abbreviations

BGC,: Biosynthetic Gene Cluster;
Bx,: Benzoxazinoids;
DIMBOA,: 2,4-dihydroxy-7-methoxy-1,4-benzoxazin-3-one;
DIMBOA-Glu,: 2,4-dihydroxy-7-methoxy-1,4-benzoxazin-3-one glucoside;
βBD,: **β**-glucosidases;
HGT,: Horizontal Gene Transfer;
HMW,: High Molecular Weight;
HTG,: Horizontally Transferred Gene;
MYA,: Million Years Ago;
MATE,: Multidrug And Toxic compound Extrusion;

