## Supplemental_information for "Horizontal gene transfer fuels metabolic innovation in the grass *Zuloagaea bulbosa*"

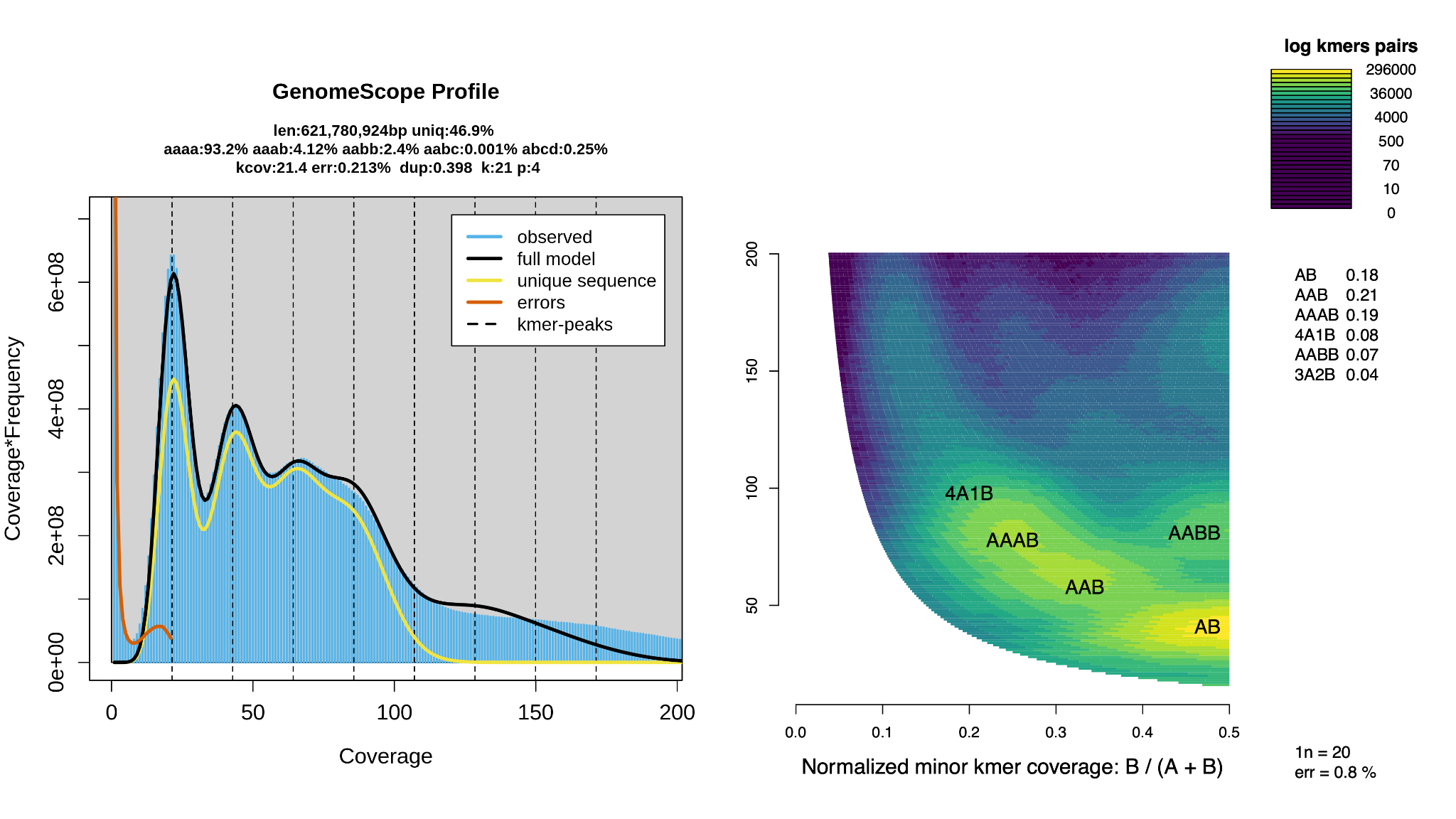

**Figure S1.** Estimation of genome size and ploidy from k-mer distribution.

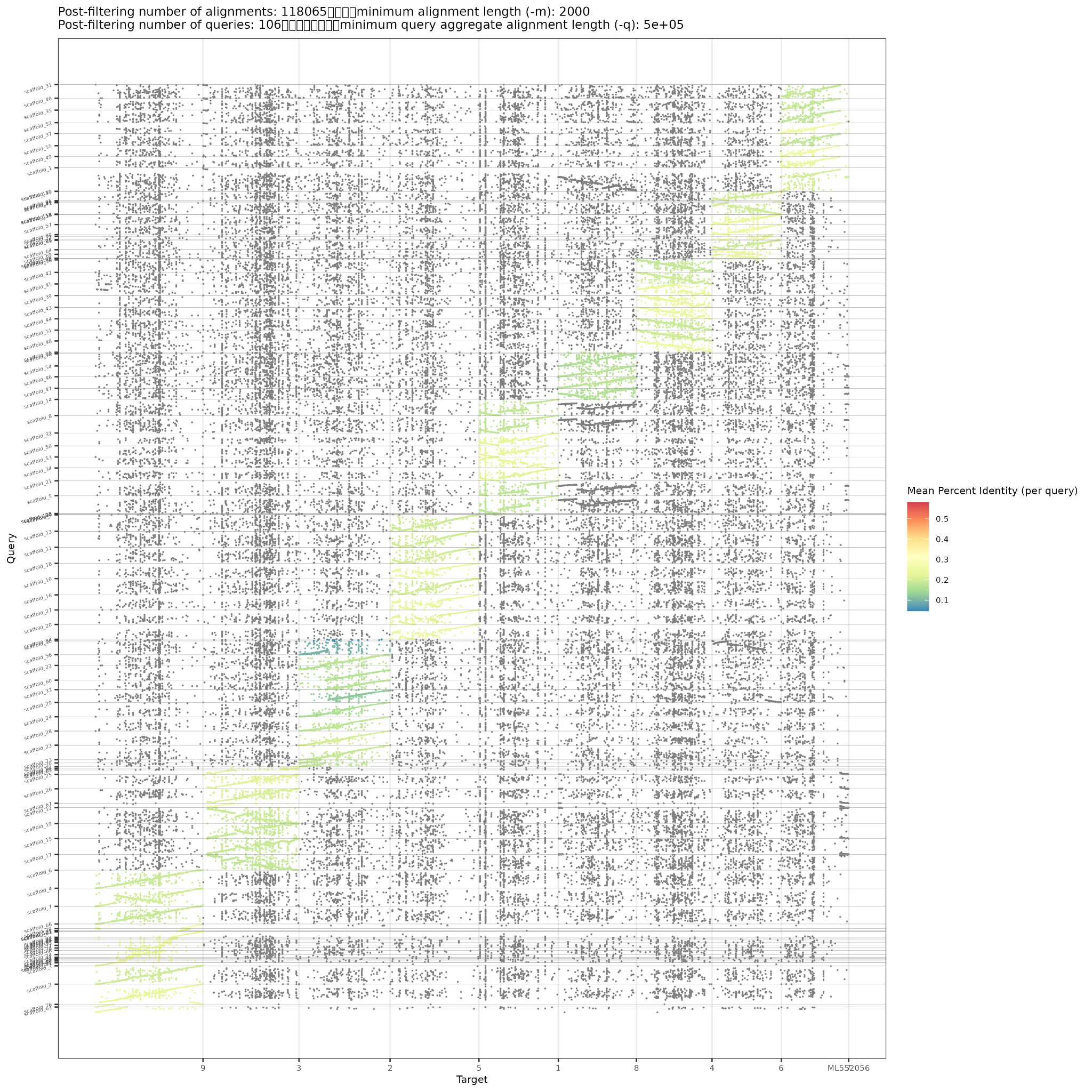

**Figure S2.** Whole-genome alignment of *Zuloagaea bulbosa* scaffolds against chromosomes of the Cenchrinae grass *Setaria viridis*, showing broad conservation of genomic collinearity between the two species.

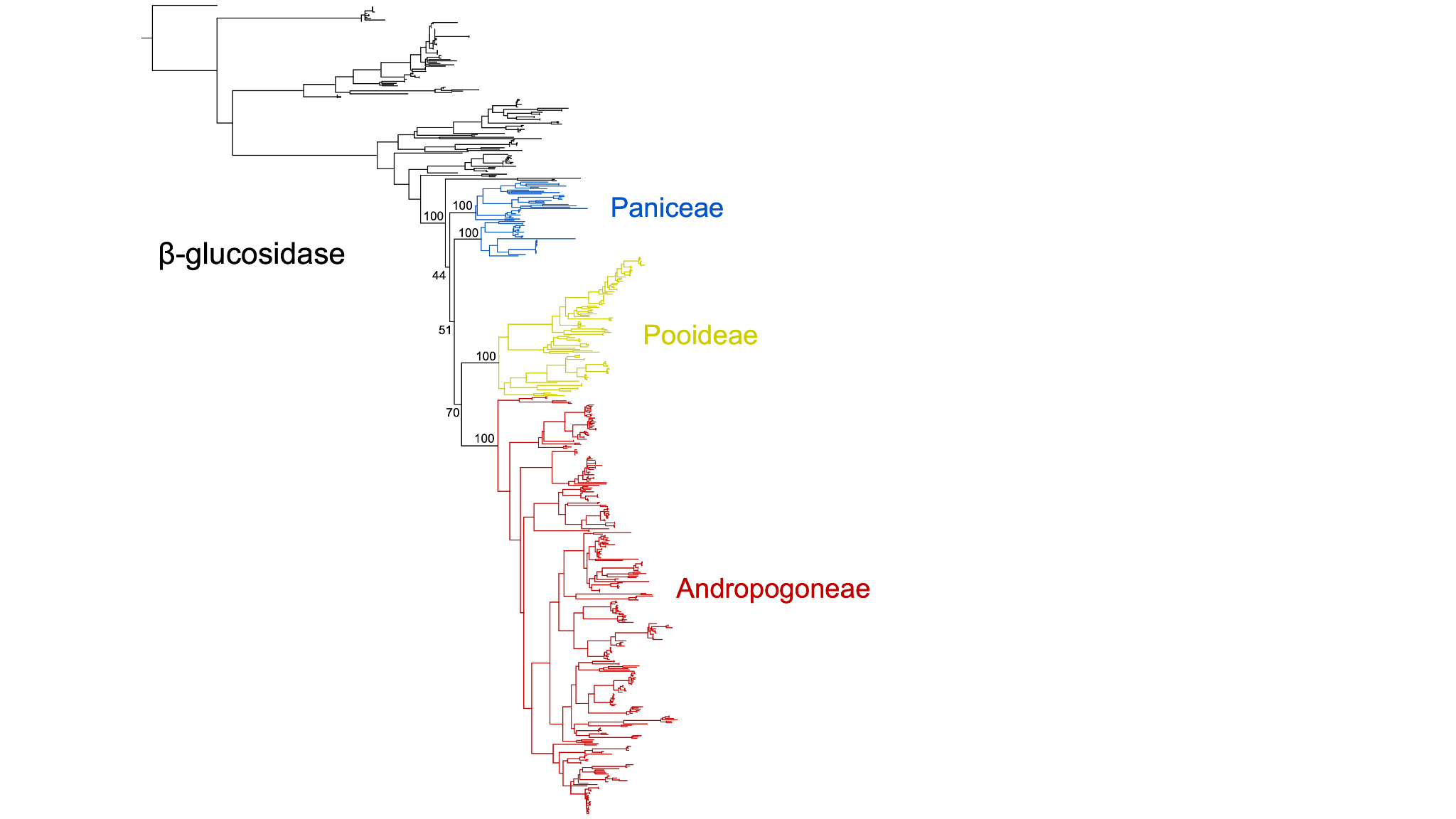

**Figure S3.** Phylogenetic tree for **𝛽**-glucosidase gene family in grasses.

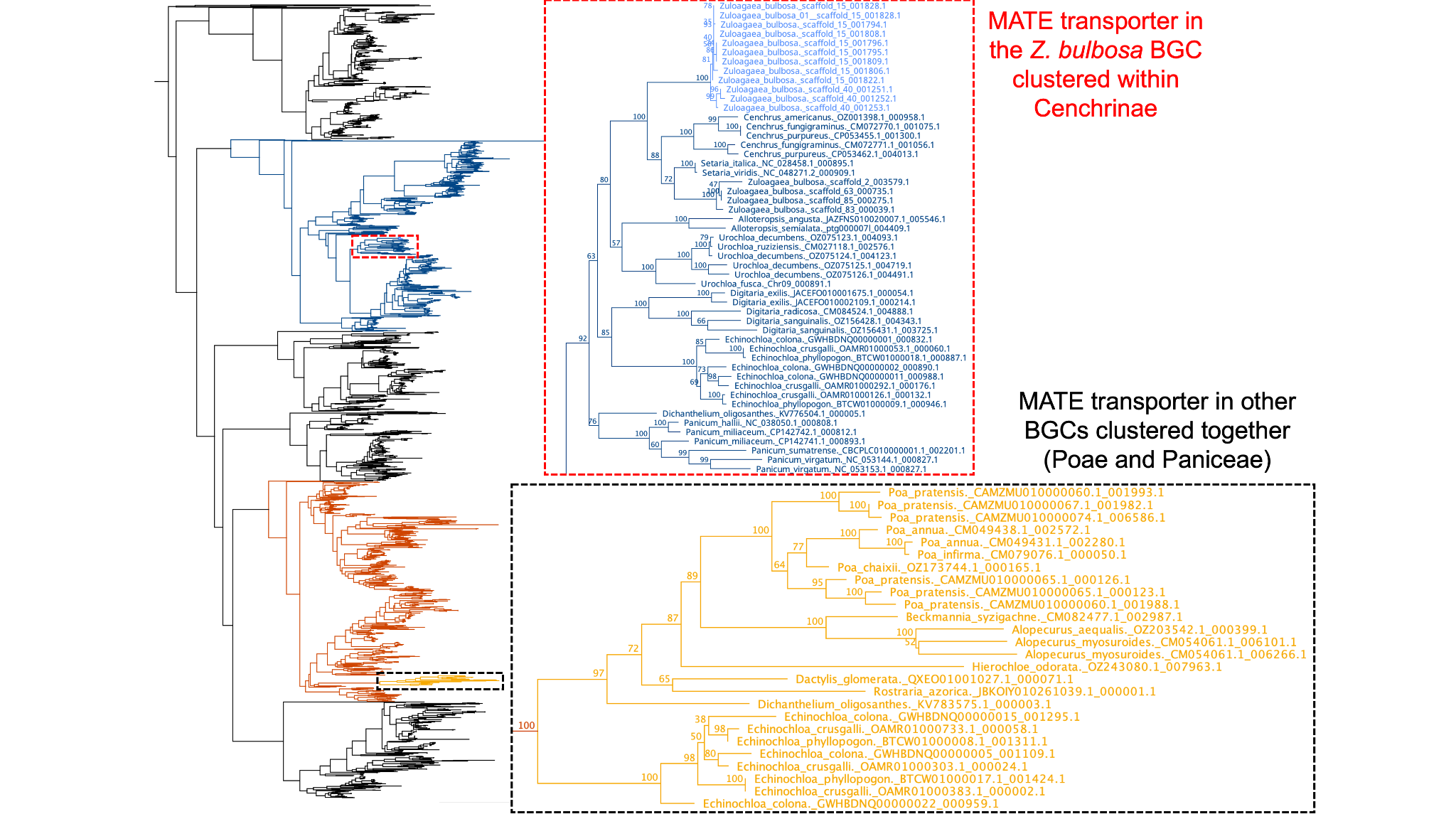

**Figure S4.** Phylogenetic tree for MATE transporter gene family in grasses.

**Dataset S1.** InterProScan functional annotation of *Z. bulbosa* predicted genes.

**Dataset S2.** Maximum-likelihood phylogenetic trees for all candidate HTGs and *Bx* genes.

**Dataset S3.** Biosynthetic gene cluster prediction from PlantiSmash. The files included in the dataset are the summary spreadsheet containing information about all clusters and individual GenBank files for individual clusters.
